# Task-evoked flexibility of intrinsic neural timescales is hierarchically organized, behaviorally relevant, and refined across two developmental stages

**DOI:** 10.64898/2026.10.21.760719

**Authors:** Linden Parkes, Valerie J. Sydnor, Amber Howell, John M. France, Andrew Westbrook, Bart Larsen

**Affiliations:** Department of Psychiatry, Brain Health Institute, Rutgers University, Piscataway, NJ, USA; Department of Psychiatry, University of Pittsburgh, Pittsburgh, PA, USA; Department of Pediatrics, University of Minnesota, Minneapolis, MN, USA; Masonic Institute for the Developing Brain, University of Minnesota, Minneapolis, MN, USA

**Keywords:** Intrinsic neural timescales, Timescale flexibility, Sensorimotor-association axis, Development, Cognition, Critical periods, fMRI

## Abstract

Cortical regions integrate temporal information over different timescales at rest, ranging from fast in sensorimotor regions to slow in the association cortex. Here, we study how far regions adapt these timescales to meet task demands, as well as how this *timescale flexibility* is organized across the cortical hierarchy, and how it develops in humans. We quantified rest-to-task changes in intrinsic neural timescales, referred to as *timescale deflections*, using fMRI data from four independent datasets totaling 1,712 participants aged 8–64 years. We found that timescales reduced under cognitive load in all four datasets, most strongly in association cortex, which partially flattened the resting-state timescale hierarchy. Individual differences in timescale deflections were organized along the sensorimotor-association (S-A) axis and tracked cognitive performance, both in adults and in youth. Next, in youth, deflections followed a two-stage developmental course: a generalized reduction throughout childhood, followed by a selective expansion in association cortex from mid-adolescence. Furthermore, the timing of these stages aligned with cortical gradients of inhibitory gene expression and intracortical myelin content, both features implicated in the opening and closing of critical periods of neural plasticity. Overall, our results demonstrate that task-evoked timescale flexibility is hierarchically organized, behaviorally and neurobiologically relevant, and refined across adolescence.

## 1 Introduction

The brain’s ability to support complex behavior depends upon integrating information across multiple timescales [1]. Reacting to a sudden change in the environment requires rapid sensorimotor processing on the order of milliseconds, whereas following a conversation or reasoning toward a goal requires integrating information over seconds, minutes, or longer. To meet this range of processing demands, cortical regions vary in terms of their temporal receptive windows (TRWs), the length of time over which a region can integrate information [1–3]. Each region’s baseline TRW is calibrated to the temporal structure of the information it preferentially processes. This produces a hierarchy of resting-state TRWs across the cortex. Primary sensory and motor regions exhibit short baseline TRWs suited to high-frequency stimulus processing, while higher-order association regions exhibit longer baseline TRWs suited to integrating information across extended time frames [4, 5]. This hierarchical organization is a fundamental feature of cortical function, observed consistently across species and measurement modalities, and its patterning is refined systematically throughout development [6–8]. However, this resting-state hierarchy reflects a single state and primarily indexes regions’ baseline capacity for temporal integration. It remains an open question how far a region’s TRW departs from this baseline with behavioral demands, how any such flexibility is organized across the cortical hierarchy, and how it emerges throughout development in humans.

Empirically, regions’ TRWs are most commonly inferred from their intrinsic neural timescales (hereafter, timescales), which quantify the rate at which a region’s local activity decorrelates from itself over time. Across species—including mice [9], macaques [10–12], and humans [13–15]—and across imaging modalities spanning electrophysiology, MEG, EEG, and fMRI, timescales reliably exhibit a hierarchical organization consistent with the cortical hierarchy of TRWs: fast timescales in primary sensorimotor cortices, progressively slower timescales in higher-order association regions. This organization is evident in the infra-slow fluctuations of the BOLD signal, where timescale gradients span the entire cerebral cortex [16, 17]. It also aligns closely with the sensorimotor-association (S-A) axis, a canonical gradient of cortical variation that captures the continuous transition from unimodal sensorimotor cortex to transmodal association cortex [18]. This hierarchical patterning of timescales is thought to arise from systematic regional differences in cortical microstructure, such as dendritic spine density, receptor and interneuron composition, and intracortical myelination [14, 15, 19, 20], which shape local circuit properties including the balance of excitation and inhibition and the strength of recurrent connectivity [19, 21].

Yet timescales are not static properties of a region. Rather, evidence suggests that regions can shift their baseline timescales as task contexts change. Evidence for this *timescale flexibility* spans diverse modalities, species, and behavioral contexts [22, 23]. In humans, MEG [24], EEG [25], ECoG [14], and fMRI [26, 27] studies report flexible speeding or slowing of timescales in different task contexts, e.g., during working memory tasks. In calcium imaging performed in rodents, timescales show relative slowing during continuous behaviors such as locomotion [25]. Thus, the task-dependent modulation of timescales may be an important feature of brain function that allows brain regions to tune their computational window within and between task contexts [12, 28, 29]. Timescales are also sensitive to changes in arousal. In human intracranial recordings, timescales slow progressively with the depth of non-rapid eye movement sleep, and the hierarchical gradient present during wakefulness is attenuated [30, 31]. Finally, recent work showed anesthesia consistently alters timescales across a diverse range of species [32]. Taken together, these cross-species and cross-modality data suggest that brain regions can be characterized by both their baseline timescale, measured at rest, and their ability to adapt their timescales in response to context and task demands, with both features being critical to understanding healthy brain function. In humans, three dimensions of this timescale flexibility remain unresolved.

The first is cognitive. If timescale flexibility represents an important computational resource in the brain, then the extent to which a region’s timescale shifts during cognitive tasks should covary with how well people perform those tasks. Direct evidence comes from human intracranial recordings, where task-evoked change in neuronal timescales tracks working memory performance [14]. However, this evidence comes from a small set of regions in a single task context, and the relationship has not been tested across the whole cortex or in relation to the hierarchy that organizes baseline timescales. Establishing this relationship would identify timescale flexibility as a source of individual variability in cognitive ability.

The second is developmental. Resting-state timescales undergo pronounced maturational change that follows the S-A axis [8, 33, 34], as do other features of cortical function, including local activity [35], as well as long-range functional and structural connectivity [36, 37]. This work establishes how baseline timescales develop, but whether the capacity to depart from that baseline in a task-dependent manner also changes with age, and whether any such change is organized along the S-A axis, remains unknown. Additionally, cognitive ability improves markedly across childhood and adolescence, with refinements to higher-order executive function occurring relatively late in adolescence [38]. This pattern suggests that context-dependent modulation of brain function may itself be maturing during mid-to-late adolescence. In turn, timescale flexibility may be one overlooked component of this developmental change, and it may mature in step with the cortical hierarchy.

The third is neurobiological. The hierarchical organization of baseline timescales along the S-A axis is thought to be underpinned by regional variation in cortical microstructure and excitatory-inhibitory (E/I) balance. E/I balance determines how long activity persists after input [19, 20], and timescales measured *in vivo* track regional variation in microstructure [15] and in the transcriptomic programs that underpin excitatory and inhibitory synaptic signaling [39–41]. Critically, *in vivo* proxies for these same properties mature along the S-A axis in humans: association cortex undergoes a selective, S-A-axis-organized shift in E/I balance across adolescence [42, 43], and intracortical myelin growth peaks late and in a graded fashion along the same axis [44]. Because these properties set the circuit dynamics that determine baseline timescales, they may also constrain how far a region’s timescale can move from that baseline, and their protracted maturation in association cortex would predict that timescale flexibility matures later in association cortex than in sensorimotor cortex. However, whether the developmental course of timescale flexibility aligns with the cortical distribution of these properties is untested. Such alignment would be consistent with the S-A critical period framework, in which the maturation of inhibitory circuitry and the accumulation of intracortical myelin open and close windows of heightened plasticity in a cascading manner along the cortical hierarchy [45–47]. In turn, these processes may facilitate the refinement of cortical microstructure that subserves timescale flexibility.

Here, we leverage four independent fMRI datasets totaling 1,712 participants [48–51] to address the following questions about the capacity of timescales to change in response to task demands: (i) how are rest-to-task timescale changes organized across the cortex, (ii) how do they vary between individuals to explain cognition, (iii) how do they emerge across development, and (iv) how does their developmental emergence align with the cortical distribution of gene expression and intracortical myelin features linked to inhibitory circuit maturation. The gaps outlined above motivate three predictions. First, if timescale flexibility is a computational resource organized by the cortical hierarchy, then its magnitude, its inter-individual variability, and its relationship to cognitive performance should each be patterned along the S-A axis. Second, if timescale flexibility matures along the cortical hierarchy, its developmental course should be heterochronous along the S-A axis rather than uniform. Third, the timing of these developmental effects should track the cortical distribution of features linked to E/I balance and intracortical myelination.

We find that timescales shorten under cognitive load in all four datasets, most strongly in association cortex, and that individual differences in this rest-to-task timescale flexibility are organized along the S-A axis and track cognitive performance. In youth, this flexibility matures in two stages, a generalized reduction through childhood followed by a selective expansion in association cortex from mid-adolescence, with a timing profile that aligns with cortical gradients of inhibitory-interneuron-related gene expression and intracortical myelin.

## 2 Results

We studied regions’ intrinsic neural timescales using fMRI data from four independent datasets, totaling 1,712 individuals aged 8–64 years (see Section 4.1). These datasets comprised two healthy adult cohorts (HCP-YA and WAND) and two developmental cohorts (HCP-D and Penn LEAD). Following our previous work [8, 52], timescales were quantified by calculating the decay lag of the autocorrelation coefficient computed separately for resting-state, movie-watching, and task-based regional fMRI time series (see Section 4.4). Here, lower values indicate faster decay of the ACF to *r* = 0, and thus faster timescales, which we interpret as shorter TRWs. For movie-watching and task-based data, we regressed out the block-design or event-related task structure from each subject’s scan before computing the ACFs (see Section 5.1 for discussion of this choice and for results without this regression applied). This choice meant that our movie-watching and task-based timescales captured fluctuations in intrinsic neural rhythms that are elicited by attentional or cognitive engagement, but that do not simply reflect the statistical structure of the scan design (e.g., the timing of the stimulus blocks). That is, the regression removed the response time-locked to the task design, not the task-related change in ongoing dynamics that we sought to measure.

### 2.1 Infra-slow timescales reduce with increasing cognitive load

Across both adults and developing youths, we found that timescales systematically reduced during tasks relative to rest (paired *t*-tests, all FDR-corrected *p <* 0.001; Fig. 1B–C), indicative of faster timescales when engaging cognitive processes. Additionally, in the HCP-YA dataset, timescales for movies also decreased relative to rest (paired *t*-test, *p <* 0.001), but not to the same extent as tasks, suggesting that timescale deflections scaled with cognitive load.

**Figure 1.**
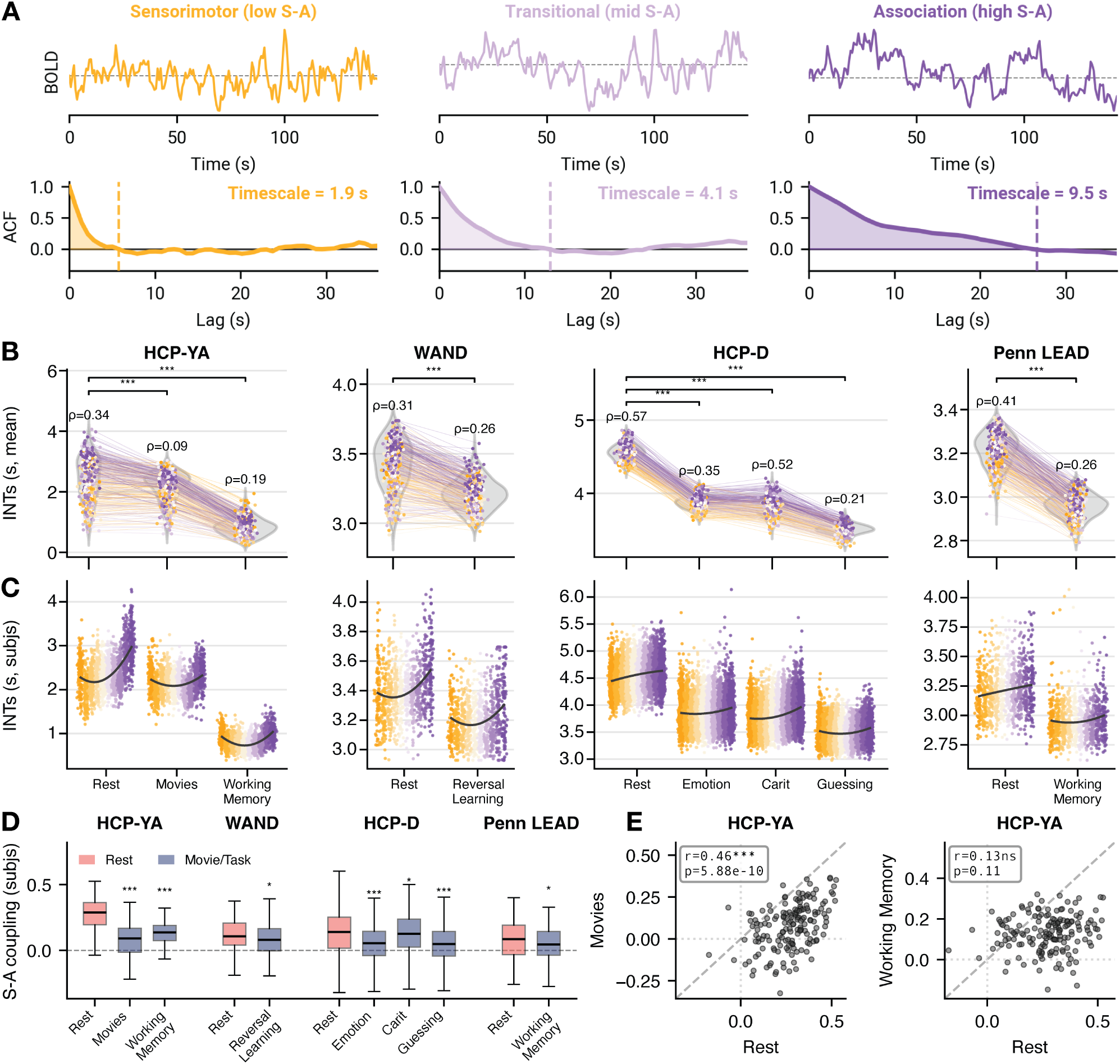
Timescales systematically reduce and decouple from the cortical hierarchy under cognitive load. (**A**) Simulated time series (top) and autocorrelation functions (ACFs; bottom) from three cortical regions spanning the sensorimotor-to-association (S-A) axis: sensorimotor (orange), transitional (light purple), and association (dark purple) cortex. The shaded area represents the timescale, computed as the area under the ACF up to the lag at which the ACF first crosses zero (dashed vertical line), multiplied by the scan repetition time. Timescales increase across the S-A axis (1.9 s *→* 4.1 s *→* 9.5 s), reflecting progressively slower autocorrelation decay in higher-order cortical regions. (**B**) Timescales reduced under cognitive load. Distributions of regional timescales, averaged over subjects, across conditions for each of the four datasets. Each point is the timescale for a single brain region averaged over subjects and colored by the S-A axis (orange = sensorimotor pole; purple = association pole). Lines connect the same region across scans, and annotated *ρ* values indicate the Spearman correlation between timescales and the S-A axis. Brackets denote paired *t*-tests across the 400 parcels, comparing each task or movie condition to rest (FDR-corrected within dataset; **\*\*\*** *p <* 0.001). (**C**) Subject-level timescale distributions grouped by S-A axis decile. Each point represents a timescale from a single subject, averaged across regions within a single decile, and the black curve traces the group-mean trend across deciles. (**D**) Timescales decouple from the S-A axis under cognitive load, assessed at the subject level. For each dataset, distributions of per-subject S-A coupling (Spearman *ρ* between a subject’s regional timescales and the S-A axis) at rest and during each condition are shown. Asterisks denote one-sided paired *t*-tests comparing each condition’s coupling to rest (Fisher-*z* transformed, FDR-corrected across all dataset *×* condition comparisons; **\*\*\*** *p <* 0.001, **\*** *p <* 0.05). (**E**) Timescales decouple differently for movies than for tasks. For HCP-YA, each subject’s S-A coupling during movies (left) and tasks (right) is plotted against their resting-state coupling; the annotated *r* values indicate the cross-subject Pearson correlation. A Steiger’s test confirmed that the rest–movie correspondence was significantly stronger than the rest–task correspondence (*z* = 3.97, *p* = 7.16 *×* 10*^−^*^5^; *n* = 163).

Note, for brevity, only the *N* -back working memory task is shown here for HCP-YA, alongside movie-watching and resting-state data averaged over the respective multiple runs (see Fig. S1 for all scans and runs). Importantly, these task-induced changes in local activity timescales, or *timescale deflections*, were not uniform across the cortex. Instead, timescale deflections were larger in association cortex, which resulted in a relative decoupling of regional timescale variation from the cortical hierarchy. Specifically, across all datasets, resting-state timescales correlated positively with the S-A axis (Fig. 1B; *ρ* = 0.34, 0.31, 0.57, 0.41 for HCP-YA, WAND, HCP-D, and Penn LEAD, respectively), consistent with previous literature. This hierarchical coupling diminished when participants transitioned from rest to watching movies (*ρ* = 0.09, HCP-YA) and performing tasks (*ρ* = 0.19, 0.26, 0.36, 0.26, where the HCP-D value is averaged over its three tasks; averaged *ρ* = 0.27 across datasets). Thus, while timescales were consistently slower in association cortex than in sensorimotor cortex at rest, this gradient partially flattened under cognitive load, driven by greater timescale flexibility in association regions relative to sensorimotor.

We assessed hierarchical decoupling at the participant level by correlating the S-A axis with participant-level timescale maps. Compared to rest, we found that S-A coupling was significantly reduced during every task and movie condition across all four datasets (one-sided paired *t*-tests, all FDR-corrected *p <* 0.05; Fig. 1D). Furthermore, in HCP-YA, we found that this participant-specific decoupling was topographically different for movies compared to the working memory task (Fig. 1E). That is, while participants’ S-A coupling was reduced under both conditions relative to rest, coupling during movies remained correlated, across subjects, with resting-state coupling (*r* = 0.46, *p* = 5.88 *×* 10*^−^*^10^), whereas coupling during working memory showed no reliable correspondence with rest (*r* = 0.13, *p* = 0.11). A Steiger’s test confirmed that this difference between movies and tasks was significant (*z* = 3.97, *p* = 7.16 *×* 10*^−^*^5^). This result suggests that movies and tasks may decouple timescales from the cortical hierarchy through distinct routes: movie-watching attenuates hierarchical coupling while preserving its between-subject variability, whereas task engagement substantially reconfigures that variability, leaving no reliable correspondence with participants’ resting baseline. Thus, individuals whose timescales are more stratified along the S-A axis at rest show stronger stratification during movies, but this resting-state hierarchical organization does not predict individuals’ task patterning.

Collectively, these results demonstrate a consistent modulatory effect of cognition on brain dynamics, in which timescales deflect from rest when the brain is driven exogenously by a diverse range of movies and cognitive tasks—with the largest deflections consistently concentrating in the association regions, indicating greater timescale flexibility relative to the rest of the cortex.

### 2.2 Timescale deflections are spatially organized and behaviorally relevant

The analyses in the previous section characterized timescales within each condition, and showed that they sped up and decoupled from the cortical hierarchy, when participants engaged in cognitive tasks. Hereafter, we change our measure of interest from timescales to the timescale deflections themselves, computed for each participant and region as the absolute difference between task and resting-state timescales (Section 4.4), and treat these deflections as an index of timescale flexibility. Thus, here, larger timescale deflections reflect greater reductions in timescales going from rest to task. This participant-level measure allowed us to ask how deflections varied between individuals across the cortical hierarchy, and whether this variation explained participants’ behavior. All analyses in this section used the working memory task from HCP-YA (*n* = 966), which yielded the largest timescale deflections relative to rest across all tasks examined (Fig. S1). Unlike Fig. 1, where resting-state timescales were averaged over all available scans, here each participant’s resting-state timescale was estimated from their 3T resting-state runs truncated to the length of the working memory run, so that rest and task estimates were matched on scan duration (see Section 4.4 for more details and Section 5 for sensitivity and robustness analyses).

We first characterized how inter-individual differences in timescale deflections were distributed across the cortical hierarchy (Fig. 2A). Both the between-subject mean and standard deviation of absolute timescale deflections increased along the S-A axis (*r* = 0.33, *p*_smash_ = 0.010; *r* = 0.34, *p*_smash_ = 0.007, respectively), indicating that association cortex not only exhibited larger deflections on average, but also showed greater inter-individual variability in the magnitude of those deflections. This result demonstrates that individual differences in timescale flexibility are disproportionately concentrated in higher-order cortex. Furthermore, this between-subject variability was highly reproducible: the spatial pattern of deflection variability estimated from the left–right (LR) acquisition was almost perfectly correlated with that estimated from the independent right–left (RL) run (*r* = 0.97; Fig. 2B). Together, these findings indicate that timescale deflections capture a stable, hierarchically organized axis of inter-individual variability in brain dynamics.

**Figure 2.**
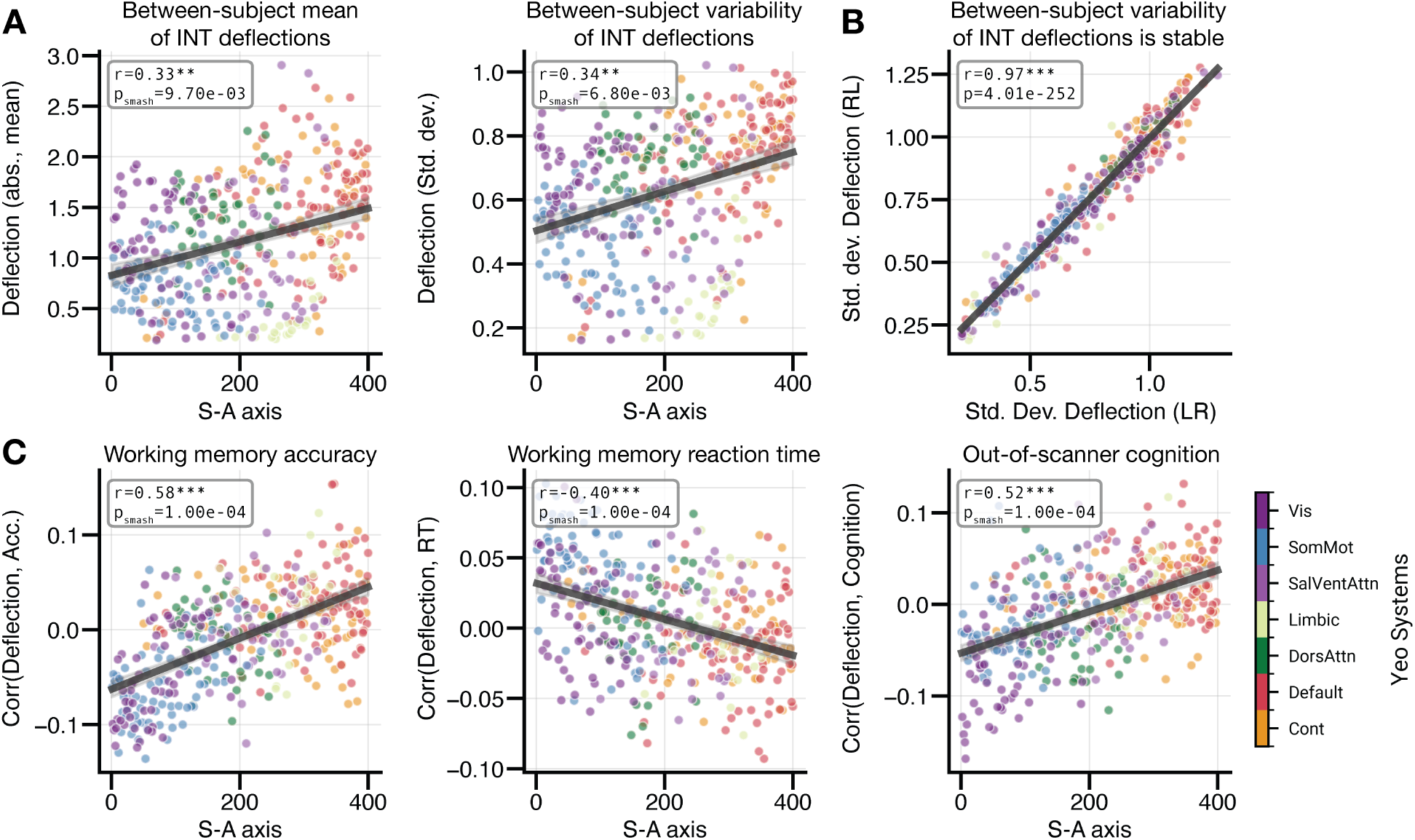
Timescale deflections are spatially organized and behaviorally relevant. All panels show HCP-YA working memory data (*n* = 966) with resting-state timescales estimated from 3T rest runs length-matched to the task run. (**A**) Between-subject mean (left) and standard deviation (right) of absolute timescale deflections plotted as a function of the S-A axis. Each point represents one brain region, colored by the canonical Yeo-7 functional systems [53]. Here, positive relationships indicate that regions higher on the S-A axis exhibit both larger and more variable timescale deflections across individuals (*r* = 0.33, *p*smash = 0.010; *r* = 0.34, *p*smash = 0.007, respectively), indicating greater flexibility. (**B**) Between-subject variability (standard deviation) of absolute timescale deflections estimated from the L–R scan acquisition plotted against the same quantity estimated from the independent R–L run. The near-perfect correlation (*r* = 0.97) indicates that the spatial pattern of timescale deflection variability was highly reproducible across independent acquisitions. Note that this panel indexes measurement reliability rather than spatial correspondence, so no spatial null was applied. (**C**) Region-wise partial correlations between absolute timescale deflections and working memory accuracy (left), working memory reaction time (middle), and an out-of-scanner measure of cognition (right), each plotted as a function of the S-A axis. Behavioral correlations were systematically organized along the S-A axis (*r* = 0.58, *p*smash *<* 1 *×* 10*^−^*^4^; *r* = *−*0.40, *p*smash *<* 1 *×* 10*^−^*^4^; *r* = 0.52, *p*smash *<* 1 *×* 10*^−^*^4^, respectively). These results indicate that larger timescale deflections in association cortex corresponded to better performance across all three measures, while larger deflections in sensorimotor cortex showed the opposite relationship.

Next, we examined whether the magnitude of timescale deflections correlated with cognitive performance, controlling for participants’ resting-state timescales, age, sex, and in-scanner motion (Fig. 2C). Region-wise correlations between timescale deflections and working memory accuracy significantly varied along the S-A axis (*r* = 0.58, *p*_smash_ *<* 1 *×* 10*^−^*^4^). In association cortex, larger deflections corresponded to greater accuracy. This relationship was reversed in sensorimotor cortex, where larger deflections corresponded to lower accuracy. This same pattern emerged for reaction time (*r* = *−*0.40, *p*_smash_ *<* 1 *×* 10*^−^*^4^): larger deflections in association cortex corresponded to faster responses, while larger deflections in sensorimotor cortex corresponded to slower responses. Thus, individuals whose fMRI-based timescales reduced more markedly (larger deflections) in higher-order association cortex—but less so in sensorimotor cortex—were both more accurate and faster on the working memory task. This result suggests that optimal cognitive performance is associated with a spatially differentiated pattern of timescale modulation, in which timescales are selectively reduced in the association cortex while sensorimotor timescales are relatively preserved. A parallel result was observed for an out-of-scanner measure of cognition (*r* = 0.52, *p*_smash_ *<* 1 *×* 10*^−^*^4^), where the same hierarchically organized pattern of behavioral correlations was present, indicating that the behavioral relevance of timescale deflections generalizes beyond in-scanner performance. Together, these results demonstrate that individual variability in the task-evoked modulation of infra-slow neural timescales is hierarchically organized and behaviorally meaningful. Note, this hierarchical organization depended on regression of the task design. Without it, neither the mean nor the variability of deflections was significantly organized along the S-A axis, and neither were the brain–behavior correlations, even though the magnitudes of those correlations were unchanged (Figs. S9 and S10; see Section 5.1).

### 2.3 Timescale deflections exhibit a two-stage developmental course throughout youth

In our previous work, we found that both the amplitude of regions’ low-frequency fluctuations [35] and their timescales [8] change systematically throughout development while participants are at rest. These developmental patterns were organized along the S-A axis: primary sensory and motor cortices tended to exhibit linear developmental decreases that onset relatively early (8-12 years), while higher-order association cortices showed non-linear trajectories that onset relatively late (18–22 years) [35]. Extending on our previous work, we next characterized the developmental course of task-based timescale modulations, including quantifying age-resolved effects to study distinct developmental phases.

Using the CARIT task in HCP-D (*n* = 552; ages 8–22 years) [50], we characterized the developmental trajectories of timescale deflections by fitting generalized additive models (GAMs; see Section 4.6)—either to regional data or to data averaged over S-A axis deciles—and examined the resulting annualized rates of change (Fig. 3). The regional heatmap of annualized change provides an overview of these trajectories across the full cortical hierarchy and age range (Fig. 3A). Qualitative assessment of this heatmap suggested that developmental refinements in timescale deflections were not uniformly expressed across age (e.g., blue and pink in Fig. 3A). Rather, reductions in timescale deflections dominated early, while deflection increases could be observed in late adolescence. To quantitatively delineate different phases of development in neural dynamics, we used change-point detection analysis, which is capable of uncovering inflection points in multivariate data. We identified that the spatial developmental pattern of regional developmental trajectories was organized into two phases: an early childhood phase that spanned from 8 to 15 years of age, and an adolescent phase that started at 15 years and continued through to 22 years.

**Figure 3.**
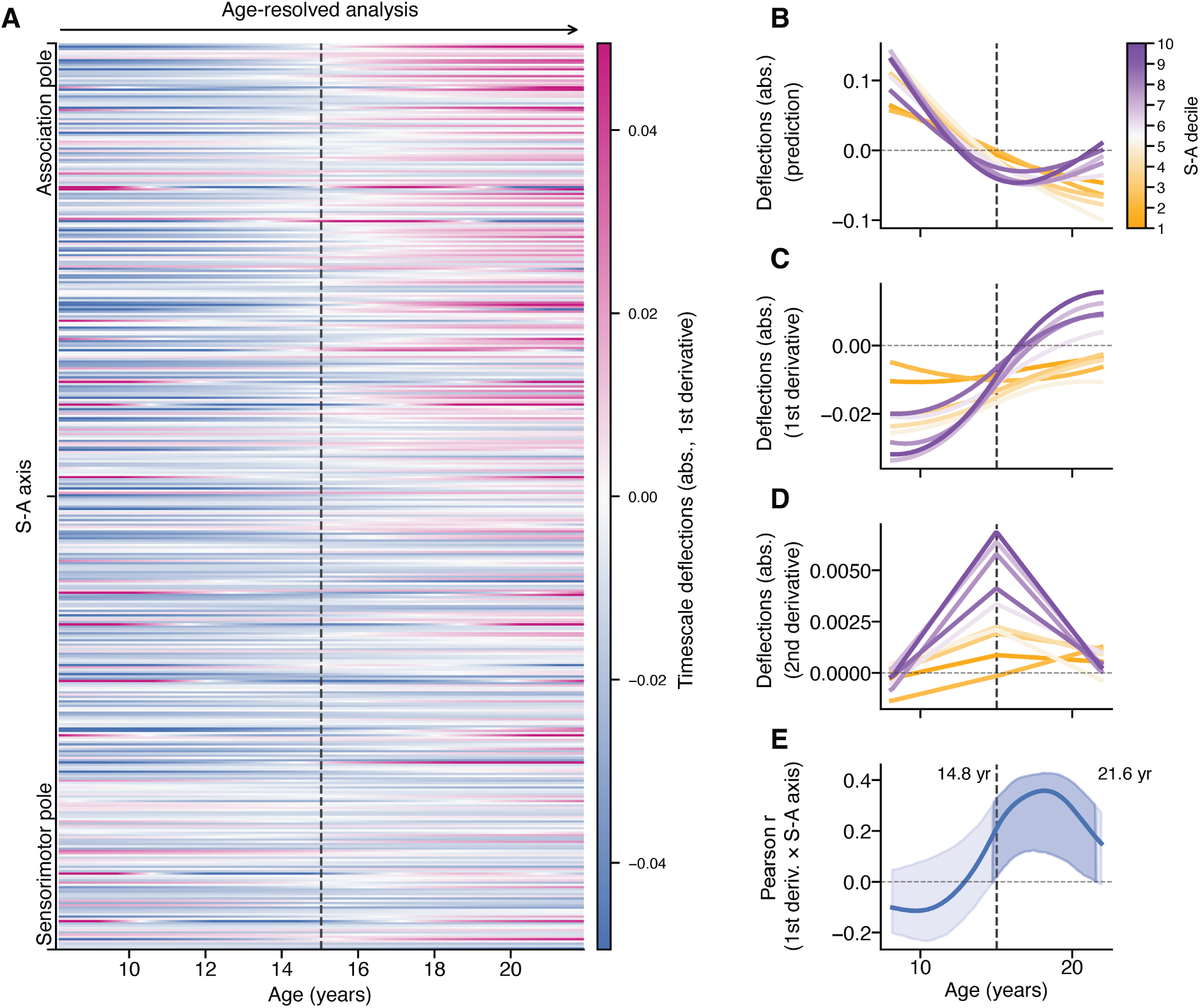
Timescale deflections exhibit a two-stage developmental course along the cortical hierarchy. (**A**) Heatmap of the annualized rate of change (1st derivative of GAM fits) in timescale deflections across development (x-axis) for each cortical region ordered by the S-A axis (y-axis; association pole at the top, sensorimotor pole at the bottom). Cool colors indicate reducing deflections, and warm colors indicate expanding deflections. The dashed vertical line marks the stage boundary at age 15. (**B**) GAM-fitted developmental trajectories of timescale deflections for each S-A axis decile (orange = sensorimotor; purple = association). Fits are mean-centered. Association deciles reduced steeply before reversing course and expanding, whereas sensorimotor deciles reduced gradually throughout development. (**C**) First temporal derivatives of the trajectories in **B**, which provide the annualized rate of change per decile. Early in development, all deciles showed negative rates, with the association cortex reducing fastest. At older ages, association deciles crossed into positive rates while sensorimotor deciles remained negative. (**D**) Second derivatives of the trajectories in **B**, which provide acceleration per decile. The developmental reduction decelerated through adolescence, most steeply in association deciles. (**E**) Pearson correlation between regional rates of change in timescale deflections and S-A axis rank, computed at monthly intervals across development. The shaded band shows the subject-level bootstrap 95% confidence interval (CI); vertical labels mark 14.8 and 21.6 years, the window over which the lower CI bound exceeded zero. Across this window, the correlation peaked at *r ≈* 0.36.

In the early phase (8–15 years), timescale deflections became smaller across the cortex. In other words, with increasing age, timescales sped up less when participants changed from being at rest to engaging in a task. In the adolescent phase, timescale deflections began to increase in association cortex while they continued to decrease in sensorimotor cortex. To confirm this pattern, we fit GAMs to data averaged within each S-A axis decile (Fig. 3B–C; see Fig. S2 for fits without mean-centering). Sensorimotor deciles showed protracted declines in timescale deflections across the full age range, whereas association deciles showed large initial declines followed by smaller increases from late adolescence. The second derivatives (Fig. 3D) showed the change in acceleration of this developmental reduction, and the peak of this change was aligned to 15 years of age. Together, these analyses describe a two-stage developmental course of timescale deflections: (i) a childhood-into-early-adolescence stage of generalized reductions across the cortex, followed by (ii) a late-adolescence stage of selective expansion in association cortex. For an extended summary of these two developmental phases see Fig. S3.

Hierarchical organization of this developmental course emerged only in mid-adolescence. To determine when, we correlated regional rates of change in timescale deflections (first derivatives from Fig. 3A) with S-A axis rank at monthly intervals, with subject-level bootstrap 95% confidence intervals around each estimate (Fig. 3E). Throughout childhood, this coupling was weak, crossing zero only at approximately 12 years. It then became reliably positive from 14.8 to 21.6 years, peaking at *r ≈* 0.36. Notably, the age at which this coupling first reached significance (14.8 years) aligned closely with the stage boundary estimated from our change-point analysis (15 years), which was estimated independently of the S-A axis. Additionally, while we focused here on timescale deflections computed for the CARIT task, the same two-stage course was evident for both the Emotion (Fig. S4) and Guessing (Fig. S5) tasks, indicating that these effects were task-general. See also Fig. S6 for an analysis of task-based timescales in the CARIT task, instead of deflections. Together, these results indicate that the cortical hierarchy governs the maturation of timescale deflections predominantly during mid-to-late adolescence, when a generalized childhood reduction gives way to selective, hierarchically organized expansion in association cortex.

Finally, as in the HCP-YA dataset, we examined whether individual differences in timescale deflections were related to task performance in HCP-D. Unlike in HCP-YA, here, regional correlations between timescale deflection magnitude and CARIT no-go accuracy were negatively organized along the S-A axis (*r* = *−*0.34, *p*_smash_ *<* 1 *×* 10*^−^*^4^; Fig. 4A). That is, larger deflections in association cortex were associated with lower accuracy, not better accuracy as in HCP-YA. Furthermore, this S-A patterning was more pronounced in the second developmental stage compared to the first (Fig. 4B). These results suggest that the behavioral relevance of timescale flexibility changes with maturation and is context specific. In youth, better inhibitory control is accompanied by smaller timescale deflections in association cortex, whereas in young adults, larger deflections are accompanied by better working memory performance.

**Figure 4.**
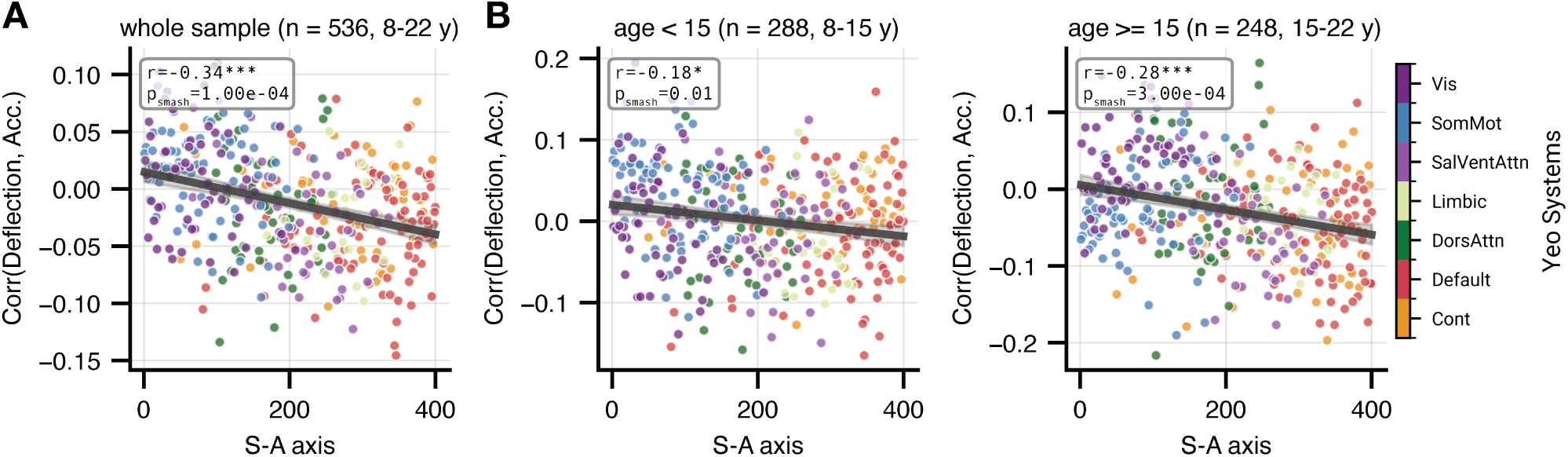
The association between timescale deflections and inhibitory control in youth tracks the S-A axis. Each point is a cortical region, colored by the Yeo-7 systems and ordered by the S-A axis (x-axis). The y-axis shows the partial correlation between regional timescale deflection magnitude and CARIT no-go accuracy, controlling for age, sex, head motion, and mean resting-state timescale. (**A**) Region-wise partial correlations for no-go accuracy computed across the whole sample (*n* = 536, 8–22 years). (**B**) Region-wise partial correlations for no-go accuracy computed separately across participants younger than 15 years (left) and participants aged 15 years and older (right). In **B**, age was still included as a covariate within each group; the 15-year split corresponds to the stage boundary in Fig. 3. \**p <* 0.05; \*\*\**p <* 0.001.

### 2.4 The two developmental stages align in sequence with cortical gradients of inhibitory and myelin-related features

The two-stage developmental trajectory outlined above raised the question of what distinguishes the childhood reduction of timescale deflections from their adolescent expansion. Because E/I balance and microstructure shape baseline timescales [15, 19] and mature along the S-A axis [42, 44], we asked whether the two stages track the cortical distribution of these properties.

To address this question, we examined three cortical maps: two gene-expression components and intracortical myelin (Fig. 5A). The two gene maps were derived from principal components analysis (PCA; see Section 4.10) applied to a curated set of *∼*2,000 brain-related genes from the Allen Human Brain Atlas (AHBA), previously used to study timescales [14]. The third map was intracortical myelin measured *in vivo* in HCP-D (see Section 4.11). The first gene component (Gene-PC1; 18.9% of variance) separated regions expressing fast-spiking interneuron machinery (e.g., parvalbumin, the Kv3 potassium channels *KCNC1*/*KCNC3*, and the Nav1.1 sodium channel *SCN1A*/*SCN1B*) from those expressing slow, tonic-modulatory signaling genes (e.g., the extrasynaptic GABA_A_ subunits *GABRA5*/*GABRB1* and the somatostatin receptor *SSTR1*). Thus, the fast-spiking pole of Gene-PC1 indexes the inhibitory machinery thought to open critical periods [54, 55]. By contrast, intracortical myelin is one of the structural brakes thought to close them [56, 57]. The third gene component (Gene-PC3; 9.0% of variance), separated regions expressing oligodendrocyte genes (e.g., the transcription factor *SOX10* and the newly-formed oligodendrocyte marker *ENPP6*) from those expressing synaptic-signaling genes (e.g., *STX1B*, *SYT7* /*SYT11*, *DLGAP1*, and the metabotropic glutamate receptor *GRM7*). Unlike Gene-PC1, which was strongly aligned with both intracortical myelin (*r* = *−*0.72, *p*_smash_ *<* 1 *×* 10*^−^*^4^) and the S-A axis (*r* = 0.68, *p*_smash_ *<* 1 *×* 10*^−^*^4^), Gene-PC3 was aligned with neither (*r* = 0.05, *p*_smash_ = 0.55; *r* = *−*0.11, *p*_smash_ = 0.21). This suggests that Gene-PC3 indexes a glial-versus-neuronal axis of cortical composition that is spatially distinct from the hierarchy, rather than a specific opener or closer of critical periods. Note, the second gene component was not associated with developmental change and is not reported further.

**Figure 5.**
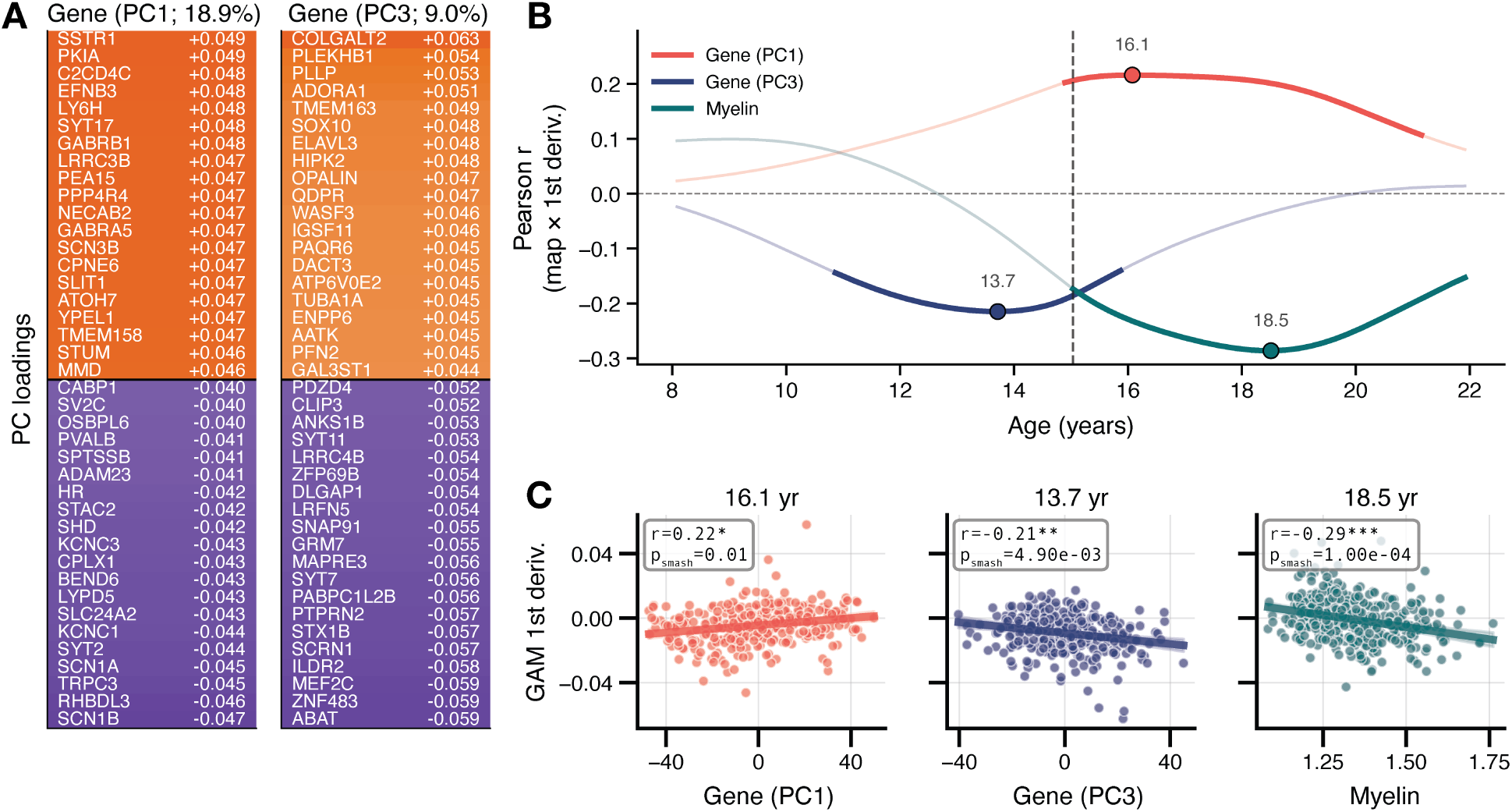
The two developmental stages of timescale deflections map onto distinct molecular organizations. (**A**) Top-loading genes at each pole of Gene-PC1 (left) and Gene-PC3 (right), with variance explained. Gene-PC1 contrasts fast-spiking interneuron machinery with slow, tonic-modulatory signaling; Gene-PC3 contrasts oligodendrocyte genes with genes related to synaptic signaling. (**B**) Pearson correlation between each of the three reference maps—Gene-PC1, Gene-PC3, and intracortical myelin measured in HCP-D—and the regional rate of developmental change in timescale deflections, as a function of age. Bold segments mark pointwise BrainSMASH nulls where *p <* 0.05, filled markers indicate the age of peak *|r|* among significant ages, and the dashed vertical line shows the stage boundary at age 15. (**C**) The regional scatter at each age peak (Gene-PC1: *r* = 0.22, *p*smash = 0.01, age 16.1; Gene-PC3: *r* = *−*0.21, *p*smash = 0.005, age 13.7; myelin: *r* = *−*0.29, *p*smash *<* 1 *×* 10*^−^*^4^, age 18.5). Each point is a cortical parcel.

We found that these maps aligned with the development of timescale deflections in sequence (Fig. 5B). Specifically, we correlated each of the above brain maps with the regional developmental rate of timescale deflections (cf. Fig. 3A) at each age. The oligodendrocyte-versus-synaptic axis (Gene-PC3) showed significant alignment with development first. In late childhood, regions rich in oligodendrocyte genes reduced their deflections fastest, and regions rich in synaptic-signaling genes reduced them least (Gene-PC3; significant across 10.9–15.9 years, peaking at 13.7 years, *r* = *−*0.21, *p*_smash_ = 0.005; Fig. 5C, middle). The fast-spiking inhibitory axis (Gene-PC1) correlated significantly with development next, starting around the transition between stages. This result suggests that regions rich in fast-spiking machinery were still reducing their deflections, whereas regions rich in tonic-modulatory signaling were changing least (Gene-PC1; 14.9–21.2 years, peaking at 16.1 years, *r* = 0.22, *p*_smash_ = 0.01; Fig. 5C, left). Finally, intracortical myelin aligned with development last and most strongly, in late adolescence. Here, heavily myelinated regions continued to reduce their deflections, whereas lightly myelinated regions expanded them (myelin; 15.0–21.9 years, peaking at 18.5 years, *r* = *−*0.29, *p*_smash_ *<* 1 *×* 10*^−^*^4^; Fig. 5C, right). These results demonstrate that each brain map marked a distinct moment in development: Gene-PC3 correlated with the childhood reduction, Gene-PC1 correlated with the transition between stages, and intracortical myelin correlated with the hierarchical divergence that followed later in adolescence.

## 3 Discussion

Here, we characterized the task-evoked modulation of infra-slow intrinsic neural timescales across four independent datasets totaling 1,712 participants. We found that timescales reduced under cognitive load across all datasets and that the magnitude of these deflections was organized along the sensorimotor-to-association (S-A) axis. Additionally, in young adults, we found that individual variability in this magnitude predicted in-scanner working memory performance and reaction time, as well as out-of-scanner cognitive ability, and in developing youths, that the same hierarchical pattern explained individual variability in response inhibition. Furthermore, in the developing brain, we found that timescale deflections followed a two-stage course, with a childhood phase of generalized reduction across the cortex followed by a late-adolescent phase of selective expansion in association cortex, and that these stages aligned in sequence with cortical patterns of gene expression and intracortical myelin. Together, these findings characterize timescale flexibility as a hierarchically organized, biologically grounded, and developmentally acquired property of the cortical hierarchy.

### Increased cognitive load leads to faster timescales in association cortex

A hierarchical organization of resting-state timescales along the S-A axis has been documented across species, modalities, and across development [8–10, 13, 14]. This hierarchy is thought to reflect systematic differences in cortical microcircuits—including regional variation in recurrent excitatory connectivity, inhibitory interneuron composition, and long-range white matter pathways—that calibrate regions’ local temporal dynamics to the demands of their specialized computational role [15, 19]. Our finding that this hierarchy is partially flattened during cognitive task engagement adds to a growing body of evidence that timescales are not static temporal-processing features but a dynamic resource [7, 24, 25, 58, 59]. By averaging over sustained task epochs across four independent datasets, we consistently observed timescale reductions under cognitive load [26, 27, 29]. This finding was robust across tasks spanning working memory, response inhibition, reinforcement learning, and naturalistic movie watching, as well as across samples ranging from childhood to adulthood. Critically, this reduction was not uniform across the cortex but caused a systematic reduction in the spatial coupling between timescales and the S-A axis, suggesting that the timescale hierarchy is selectively reorganized rather than globally compressed under cognitive demand [17].

In the HCP-YA dataset, where both movie-watching and task data were available, timescale deflections scaled with cognitive load, providing further evidence that infra-slow timescale modulation is graded by the degree of cognitive engagement rather than reflecting a binary switch between rest and task. More broadly, this selective, state-dependent reorganization fits an emerging principle whereby dynamic properties of cortical activity—beyond baseline timescales—are organized along the S-A axis and reconfigure with cognitive state. Thus, the cortical hierarchy may be a common scaffold along which the brain reallocates distinct dynamic resources as processing demands change.

Our findings of task-evoked changes in timescales builds upon an emerging body of work demonstrating that timescales flexibly adjust according to context and state. Using intracranial recordings in humans, Krause *et al.* estimated timescales from local field potential data and tracked them across sleep stages [31]. Timescales slowed progressively with the depth of non-rapid eye movement sleep, and the hierarchical organization present during wakefulness diminished. Thus, cognitive load and sleep appear to move timescales in opposite directions—faster under task, slower under sleep—yet both attenuate the cortical hierarchy. Furthermore, cognition and sleep also appear to attenuate the hierarchy from opposite ends. During sleep, hierarchical flattening was driven by a selective slowing of sensorimotor timescales toward association values [31]. By contrast, in our data, it was driven by a speeding up of association timescales toward sensorimotor values (Fig. 2A). Anesthesia, however, shows a different set of results. Deep anesthesia speeds up infra-slow fMRI timescales, most strongly where baseline timescales are slowest [32], resembling the effect of task engagement in our data. This effect may also depend on dose: propofol sedation slows fMRI timescales, whereas surgical-level anesthesia speeds them up [60]. Because propofol acts largely by potentiating GABA_A_-mediated inhibition [32], this dose dependence suggests that inhibition can either slow or speed infra-slow timescales. This interpretation is consistent with evidence that focal inhibition can strengthen slow cortical rhythms [61, 62]. Taken together, these comparisons indicate that different brain states depart from the resting-state hierarchy in distinct, spatially patterned ways. Whether these different states lie along a common axis, or share underlying mechanisms, warrants further investigation.

### Hierarchically-organized timescale flexibility predicts cognitive performance

We found that the S-A axis not only organizes the magnitude of timescale deflections, it organizes between-subject variability in timescale deflections and explained cognitive performance. Association cortex exhibited not only larger mean deflections but also greater inter-individual spread in deflection magnitude. Behaviorally, in young adults, individuals whose timescale deflections were largest in association cortex and relatively preserved in sensorimotor cortex were both more accurate and faster on the working memory task, and showed higher out-of-scanner cognition. To our knowledge, the most direct prior evidence for the behavioral relevance of task-evoked timescale change comes from intracranial recordings, in which prefrontal deflections during working memory maintenance predicted individuals’ task accuracy [14]. Our findings extend this result from a small subset of regions to the whole cortex, where the behavioral relevance of timescale deflections followed the S-A axis, and from electrophysiological to infra-slow fMRI dynamics in large samples. Thus, in adulthood, individuals who most effectively shift their infra-slow dynamics in higher-order association cortex are better positioned to respond when cognitive demand is highest, yielding advantages in both response accuracy and speed.

In youth, however, we found that the direction of this hierarchical relationship was reversed. Specifically, in HCP-D, larger timescale deflections in association cortex were accompanied by poorer inhibitory control on the CARIT task (Fig. 4). Furthermore, this negative hierarchical patterning was more pronounced after mid-adolescence (Fig. 4B), when timescale deflections began to expand selectively in association cortex. These results demonstrate that the behavioral relevance of timescale flexibility may be context specific, with both larger and smaller deflections in association cortex linking to improved task performance depending on the needs of the behavioral domain. Thus, our findings suggest that it may be the capacity for larger deflections in association cortex that supports behavior, not the direction itself. In turn, this implies that greater timescale flexibility in association cortex confers a functional advantage not necessarily through greater deflections, but through greater precision of timescale tuning to a specific task context.

### Timescale deflections develop through two distinct stages along the cortical hierarchy

Confirming our prediction that timescale flexibility develops heterochronously along the cortical hierarchy, we observed a two-stage developmental pattern: a childhood phase of broad reductions in timescale deflections that was followed by a mid-to-late adolescent phase of selective expansion in higher-order association cortex. Notably, these stages map onto well-established models of neurodevelopment. The first stage coincides with several key global developmental processes that are prevalent throughout childhood and early adolescence. For example, synaptic pruning proceeds throughout this window, beginning earlier in primary sensory cortex than in prefrontal cortex [63], where spine density continues to decline into the third decade [64]. Inhibitory interneuron circuitry matures in parallel. In the human prefrontal cortex, markers of fast-spiking interneurons rise across childhood and peak in the teenage years [65]. Theoretical work suggests that the maturation of inhibition suppresses spontaneous activity to a greater extent than it suppresses stimulus-driven activity [66]. In turn, this is thought to shift the cortex toward an externally driven operating mode and enhances the signal-to-noise ratio of cortical processing [18]. Consistent with this account, the genetic axis separating oligodendrocyte genes from synaptic-signaling genes (Gene-PC3) coupled to developmental change predominantly during this first stage, peaking at 13.7 years, with regions rich in oligodendrocyte machinery reducing their timescale deflections fastest. Because Gene-PC3 was aligned with neither the S-A axis nor intracortical myelin, this early coupling is not attributable to the hierarchy. Instead, it may reflect regional differences in the balance of oligodendrocyte-lineage and neuronal synaptic components during the period when broad synaptic refinement is underway.

By contrast, the second stage—the selective expansion of timescale deflections in association cortex—emerged in mid-to-late adolescence, becoming robustly organized along the S-A axis between approximately 14.8 and 21.6 years. This timing aligns with the protracted, late maturation of transmodal association cortex that distinguishes human neurodevelopment [18]. *In vivo*, association cortex undergoes a selective, S-A-axis-organized shift in its E/I balance across adolescence [42], accompanied by graded, late-arriving intracortical myelination [44]. Consistent with these mechanisms, recent cross-species work has shown that resting-state timescales in frontal and parietal association cortex are refined throughout adolescence in concert with E/I and deep-layer myelination changes [67]. Our results extend this account from resting-state dynamics to their task-evoked flexibility, suggesting that the same protracted maturation that refines association-cortex timescales at rest may also elaborate the capacity of these systems for context-dependent modulation.

These effects complement those we previously reported for the amplitude of low-frequency fluctuations [35]. In that work, the alignment between age-resolved developmental change and the S-A axis was strongest in childhood and early adolescence, peaking near age 15 and then weakening through late adolescence. The hierarchical organization of timescale-deflection development followed the opposite trajectory over the same period: it was absent in childhood and emerged only in mid-adolescence, becoming reliable after age 14.8. The two, therefore, overlap in time but move in opposite directions. As the cortical hierarchy’s relationship with resting-state development diminishes from its early-adolescent peak, its connection to the task-evoked flexibility of timescales rises and plateaus. Thus, rather than maturing in separate windows, the two succeed one another within a shared mid-adolescent window, as hierarchical organization shifts from resting-state activity to its task-evoked modulation. Consequently, the cortical hierarchy appears to shift from organizing regions’ intrinsic activity at rest to organizing how flexibly that activity can be reconfigured under cognitive demand. This crossover coincides with the canonical stabilization of executive function in mid-to-late adolescence [38], raising the possibility that the hierarchical elaboration of task-evoked timescale flexibility helps consolidate higher-order cognition as the earlier, resting-state mode of hierarchical development resolves.

### Timescale flexibility matures in an order consistent with a critical-period cascade

Prior work suggests that critical periods of heightened neural plasticity open as parvalbumin-expressing interneurons mature and shift E/I balance [54, 55, 68], and that these critical-period windows close as perineuronal nets, oligodendrocytes, and intracortical myelin accumulate across development [57, 69–71]. Importantly, these windows close earlier in sensory cortex than in association cortex [45, 72–75], and we have previously proposed that this cascade partly drives the S-A axis of human cortical development, with adolescence forming a late critical period in association cortex that coincides with the refinement of executive function [46, 47, 76]. The order of the couplings we observed is consistent with this account. The fast-spiking pole of Gene-PC1, which contains inhibitory machinery thought to open critical periods, coupled to development from around the stage transition. Regions at the fast-spiking pole, predominantly in sensorimotor cortex, continued to reduce their deflections, while association regions changed least and then expanded. The adolescent expansion was therefore concentrated in association regions that are relatively sparse in this machinery in adulthood, which fits with evidence that inhibitory circuitry in association cortex matures late [65]. Intracortical myelin, a principal brake on plasticity [56, 57], coupled later and for longer, peaking at 18.5 years, when heavily myelinated sensorimotor regions continued to reduce their deflections while lightly myelinated association regions expanded. Thus, the order of developmental couplings in our results is consistent with the order in which critical-period windows are thought to open and then close.

### Limitations

Several limitations of the present study warrant acknowledgment. First, all findings are derived from fMRI, which is constrained by its temporal resolution and the indirect hemodynamic coupling between neural activity and the BOLD signal. As such, the infra-slow timescales we measured here represent macroscale aggregates of many underlying neural processes, and findings based on BOLD fluctuations do not always align with those from methods that capture faster neural dynamics [22, 23, 32, 77]. For example, timescales measured with intracranial electrophysiology lengthened in prefrontal cortex during working memory maintenance [14], whereas we observed infra-slow timescales shortening under working memory load. Whether such differences reflect distinct processes operating at different frequencies, or differences in what different modalities measure, remains an open question that will require multi-modal validation, including against the spiking-level timescales documented in non-human primates [10–12]. Second, the task regression approach we applied—removing block-design or event-related task structure from the BOLD time series before computing ACFs—is a necessary step for estimating intrinsic dynamics during task engagement, but it cannot fully disentangle intrinsic neural fluctuations from residual stimulus-evoked variance, particularly in densely-sampled event-related designs [78]. Third, our developmental analyses were cross-sectional; the two-stage trajectory we describe is inferred from age-related differences across participants rather than tracked within individuals, and longitudinal studies are needed to confirm the timing, shape, and inter-individual variability of this trajectory. Fourth, our molecular analyses relate static, group-level maps of gene expression and intracortical myelin to the rate of change of a functional phenotype. The correspondence we report is therefore between spatial patterns and developmental timing, and does not establish that these molecules cause the flexibility of intrinsic timescales. Gene-PC1 and the myelin map are both strongly aligned with the cortical hierarchy, and with each other, so their ordering rests on the temporal separation of their couplings rather than on their spatial independence. Gene-PC3, by contrast, was correlated with neither, so its earlier coupling is not attributable to the hierarchy. Fifth, regional fMRI timescales cannot distinguish a fixed circuit whose dynamics are modulated between rest and task from the differential engagement of local subpopulations with distinct intrinsic timescales. Either would shift the regional timescale, and we leave disentangling these underlying mechanisms to future work.

### Conclusions

In this work, we characterized how intrinsic neural timescales reconfigure from rest to task, and showed that the flexibility of infra-slow cortical dynamics was hierarchically organized. The magnitude of this flexibility was graded along the sensorimotor–association axis and tracked how well individuals performed during cognitive tasks. We also found that this flexibility may be acquired in two developmental phases: a broad childhood reduction, followed by a mid-adolescent reorganization tightly aligned to the cortical hierarchy, with each phase tracking distinct molecular features of the cortex. In this later phase, hierarchical organization shifted from the resting-state dynamics that dominate earlier development to task-evoked modulation, coinciding with the consolidation of higher-order cognition. Together, these findings establish task-evoked timescale flexibility as a distinct, developmentally acquired axis of cortical function and position its mid-adolescent, hierarchically organized emergence as a candidate substrate for the maturation of complex behavior.

## 4 Materials and Methods

### 4.1 Participants

We analyzed neuroimaging data from four independent datasets, totaling 1,712 participants spanning ages 8.1 to 63.7 years. These included 966 adults from the Human Connectome Project Young Adult dataset (HCP-YA; ages 22.0–37.0, *M* = 28.7, *SD* = 3.7) [48], 95 participants from the Welsh Advanced Neuroimaging Database (WAND; ages 18.7–63.7, *M* = 28.7, *SD* = 9.0) [49], 552 youth from the Human Connectome Project Development dataset (HCP-D; ages 8.1–21.9, *M* = 14.9, *SD* = 3.9) [50], and 99 youth from the Penn LEAD study (ages 8.6–16.5, *M* = 12.6, *SD* = 2.0; first session only) [51]. Across the combined sample, 916 participants (53.5%) were female. All datasets received ethical approval from their respective institutional review boards, and all participants—or their legal guardians for minors—provided written informed consent prior to participation. Analyses involving HCP-YA movie-watching data were restricted to the 163 HCP-YA participants with 7T sessions (ages 22.0–36.0, *M* = 29.5, *SD* = 3.3; 98 female, 60.1%), as movies were acquired only at 7T.

Owing to known confounds between head motion and development neuroimaging [79–81], subjects were excluded from analysis if they exceeded a mean framewise displacement (FD) threshold of 0.2 mm in any included functional scan. This criterion was applied to HCP-D, yielding the 552 subjects noted above (mean FD = 0.128 *±* 0.033 mm; range 0.033–0.200 mm), and to WAND, which reduced from 160 to the 95 subjects noted above (mean FD = 0.131 *±* 0.030 mm; range 0.062–0.183 mm). For HCP-YA, no subjects were excluded based on motion. Instead, relative RMS displacement was extracted for use as a nuisance covariate (mean relative RMS = 0.088 *±* 0.031 mm; *n* = 966). Similarly, we retained the full sample for Penn LEAD to preserve sample size (mean FD = 0.352 *±* 0.354 mm; range 0.075–2.497 mm). For all datasets, motion was included as a nuisance covariate where between-subjects analyses were performed.

### 4.2 MRI data acquisition

Full MRI acquisition parameters for each dataset—including scanner hardware, sequence design, and acquisition structure—are detailed in the original protocol and data-descriptor publications, to which we refer the reader: HCP-YA [48, 82, 83], HCP-D [50, 84], WAND [49], and Penn LEAD [51].

### 4.3 MRI data processing

All datasets were processed to produce a common set of regional BOLD time series parcellated according to the Schaefer-400 atlas [85]. Pre-processing proceeded along one of two paths: (i) minimally pre-processed data was sourced from the HCP-YA and HCP-D consortiums and additional steps outlined below were applied, and (ii) fMRIPrep and XCP-D were used in conjunction to process WAND and Penn LEAD. We describe these processing streams in more detail below.

#### 4.3.1 Processing the Human Connectome Project

The HCP datasets [48, 48, 50, 82–84] consist of multi-modal neuroimaging data from young adult (HCP-YA) and developing youths (HCP-D).

##### Functional magnetic resonance imaging (fMRI)

For both HCP-YA and HCP-D, minimally preprocessed fMRI data were parcellated using the 400-region variant of the Schaefer parcellation [85]. Concretely, this involved using, for example, rfMRI_REST1_LR_Atlas_MSMAll_hp2000_clean.dtseries.nii for resting-state data and tfMRI_WM_LR_Atlas_MSMAll.dtseries.nii for task data from the S1200 release for HCP-YA. Thus, these data differed in terms of their minimal preprocessing, with the former including ICA-FIX denoising and high-pass filtering not applied to the latter [86]. For HCP-D, minimal processing was consistent across rest and task, and we used files labeled [scan]_Atlas_MSMAll_hp0_clean.dtseries.nii. For both datasets, additional nuisance regression was performed that comprised the global signal (GS).

##### Behavior

For HCP-YA, we extracted in-scanner accuracy and reaction time from the *N* -back working memory as well as a composite measure of cognition measured outside the scanner (CogTotalComp_Unadj). We correlated these variables against timescale deflections, controlling for age, sex, in-scanner motion, and resting-state timescales. For HCP-D, we extracted in-scanner performance on the CARIT go/no-go task. Specifically, CARIT no-go accuracy was computed as the proportion of no-go trials on which participants correctly withheld a response (i.e., one minus the no-go false-alarm rate), pooled across previously rewarded and previously punished no-go stimuli.

#### 4.3.2 Processing the WAND and Penn LEAD datasets

Raw, bids-compliant fMRI data from the WAND [49] and Penn LEAD [51] datasets were both processed using *fMRIprep* [87, 88] and *XCP-D* [89]. Full details of these processing steps can be found in the *boilerplate* content included below in Section 5.1.3. As above, regional fMRI time series were extracted using the 400-region variant of the Schaefer parcellation [85].

### 4.4 Intrinsic neural timescales

We quantified each region’s intrinsic neural timescale (timescale) by calculating the integral to zero-crossing of the autocorrelation function (ACF) [8, 52, 90, 91]. For each subject, scan, and Schaefer-400 parcel, we computed the ACF of the regional BOLD time series and defined the timescale as the area under the ACF from lag 0 to the lag at which the ACF first crossed zero, multiplied by the scan’s repetition time to express the timescale in seconds. Lower values indicate faster decay of the autocorrelation, and thus faster timescales or shorter temporal receptive windows. For task-based and movie-watching data, we first regressed the task structure—the block or event-related design for task runs, and the alternation between movie clips and interleaved rest periods for movie runs—from each subject’s regional time series [78, 92] and computed the ACF from the residuals. Note that this step was not possible for WAND, for which the task design was not available; WAND task timescales were therefore estimated from unregressed time series. Regressing out task structure meant that timescale estimates reflected intrinsic neural dynamics during cognitive engagement rather than stimulus-locked responses. See Section 5.1 for a close examination of this choice, and its impact on our key results. To capture task-evoked modulation of intrinsic dynamics, we defined a timescale deflection per region per subject as the difference between that subject’s resting-state and task-based or movie-watching timescales (task/movies minus rest). With the exception of Fig. 1, all analyses used the absolute value of this deflection quantity, *|*Δtimescale*|*, for which larger values index greater task- or movie-evoked speeding up of timescales.

For the HCP-YA working memory analyses in Fig. 2 and the HCP-D CARIT analyses in Figs. 3 and 5, each subject’s resting-state timescale was estimated from their resting-state runs truncated from scan onset to the duration of the corresponding task run, so that rest and task timescales were matched on scan length [93]. For HCP-YA this differs from Fig. 1, in which resting-state timescales were averaged across all available 3T and 7T resting-state scans at full length.

### 4.5 Sensorimotor–association axis

Throughout, we organized cortical regions along the sensorimotor–association (S-A) axis, a dominant axis of cortical organization running from primary sensory and motor cortex to transmodal association cortex [18, 35]. We used the S-A axis map projected to the Schaefer-400 (7-network) parcellation, and rank-transformed its values to yield an integer S-A rank from 1 at the sensorimotor pole to 400 at the association pole. We used this rank primarily in two ways. For region-level analyses, we correlated cortical maps directly against the S-A rank. For decile-level analyses, we sorted regions by ascending S-A rank and averaged them into ten equally sized bins of 40 regions, from the sensorimotor pole (decile 1) to the association pole (decile 10).

### 4.6 Generalized Additive Models

To characterize developmental trajectories of timescale deflections, we fit generalized additive models (GAMs) using the pygam library. GAMs capture non-linear predictor–outcome relationships through penalized smooth basis functions, making them well-suited to modeling development. Here, our outcome variables were the absolute timescale deflections, *|*Δtimescale*|*. We fit a separate GAM for each Schaefer-400 parcel, and, in a parallel analysis, for deflections averaged within S-A axis deciles. Each model took the form 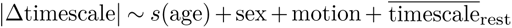. Because each deflection is defined relative to rest, its magnitude depends partly on a participant’s resting-state baseline; participants with longer resting-state timescales have more room for their timescales to reduce during a task. Resting-state timescales also change across development [8, 33, 35]. As such, we included each participant’s whole-brain mean resting-state timescale as a covariate, so that age effects on deflections reflect changes in timescale flexibility beyond those attributable to developmental shifts in the global resting baseline. The age term *s*(age) was a penalized cubic B-spline with five basis functions, with the smoothing penalty *λ* selected separately for each region by generalized cross-validation (GCV). Sex, mean framewise displacement, and each subject’s whole-brain mean resting-state timescale were entered as linear terms, and each series was mean-centered before fitting. To recover the age trajectory, we evaluated each fitted model at 200 equally spaced ages spanning the observed range, holding the covariates at their sample means. The rate of developmental change was estimated using the first derivative of this curve, approximated by finite differences between adjacent ages. For the age-resolved hierarchical analysis (Fig. 3E), we computed the Pearson correlation between regional rates of change and S-A axis rank at each age. We estimated 95% confidence intervals by resampling participants with replacement over 5000 iterations, refitting all GAMs and recomputing the correlation within each iteration.

### 4.7 Spatial null model

To assess the statistical significance of spatial correlations between cortical maps we used BrainSMASH (Brain Surrogate Maps with Autocorrelated Spatial Heterogeneity) [94]. BrainSMASH generates surrogate maps that preserve the empirical spatial autocorrelation of an input map by fitting the relationship between map values and pairwise inter-region distances (the empirical variogram), then reproducing this variogram through randomization and smoothing. Distances were surface geodesic distances between parcel centroid vertices, defined within each hemisphere; surrogates were generated per hemisphere and concatenated. For any given comparison between two brain maps, we generated surrogates for one of the two maps 10,000 times and computed its correlation with the other, unpermuted map. Then, we compared the observed correlation to the surrogate correlation-null to obtain a non-parametric p-value, denoted *p*_smash_, as the proportion of surrogates whose correlation was at least as extreme as the observed value. This test was two-tailed—comparing surrogate and observed *|r|*—for all analyses.

### 4.8 Brain–behavior associations

To relate timescale deflections to behavior, we computed, separately for each Schaefer-400 parcel, the Pearson partial correlation across subjects between the absolute timescale deflection (*|*Δtimescale*|*) and each behavioral measure (see Section 4.3.1). In HCP-YA, covariates were age, sex, in-scanner motion (relative RMS displacement) during both the resting-state and working memory runs, and each subject’s mean resting-state timescale across parcels (*n* = 966). In HCP-D, covariates were age, sex, mean framewise displacement, and each subject’s mean resting-state timescale across parcels; these analyses included the 536 of 552 participants with CARIT performance data. This procedure yielded, for each behavioral measure, a cortical map of regional brain–behavior correlations. We quantified the hierarchical organization of each map as its correlation with S-A axis rank, and assessed significance against 10,000 BrainSMASH surrogates of the S-A axis (Section 4.7). For the age-split analysis in HCP-D (Fig. 4B), we repeated this procedure separately in participants younger than 15 years (*n* = 288) and in those aged 15 years and older (*n* = 248), retaining age as a covariate within each group.

### 4.9 Two-stage developmental model

As discussed in the main text, the regional rates of change in timescale deflections (*|*Δtimescale*|*) transitioned from a broad reduction to a hierarchically patterned mixture during adolescence (Fig. 3), which we modeled as a two-stage developmental process. To locate the boundary between stages, we applied change-point detection to the age-resolved first derivative of the regional deflection fits using the ruptures library [95], configured to return a single change point, which it placed at 15 years (subject bootstrap 95% CI 14.8–15.1 years). We then defined an earlier window (stage 1; ages 8–15) and a later window (stage 2; ages 15–22). For each Schaefer-400 parcel, we integrated the fitted GAM first derivative across each window to obtain the net change in *|*Δtimescale*|* over that stage. This integral equals the difference in fitted values at the window endpoints. Dividing each net change by its window duration yielded an annualized rate of change, giving a stage-1 map and a stage-2 map across the cortex (Fig. S3). The 15-year boundary was corroborated by an independent criterion: the age at which the correlation between regional rates of change and the S-A axis first became reliably positive (14.8 years; Fig. 3E), which fell within 0.3 years of it.

### 4.10 Gene expression from the Allen Human Brain Atlas

To characterize the molecular organization of the two developmental stages, we related cortical gene expression to the regional rate of developmental change in timescale deflections. We analyzed microarray data from the Allen Human Brain Atlas (AHBA), the full details of which can be found elsewhere [96]. Briefly, the AHBA includes normalized genome-wide microarray data (58,692 probes) for 3702 tissue samples taken from the cortex and subcortex of six post-mortem adult brains. We have used these data previously in combination with *in vivo* neuroimaging to bridge across multiple scales of neuroscience [52, 97].

Using the abagen toolbox (v0.1.3) with default parameters [98], we extracted parcellated whole-brain maps of gene expression following an established processing pipeline [99]. Microarray probes were reannotated [99], and probes not matched to a valid Entrez ID were discarded. Probes with expression intensity below background in at least 50% of samples across donors were removed [100], and, where multiple probes indexed the same gene, we retained the probe with the most consistent pattern of regional variation across donors (i.e., differential stability) [101]. Tissue samples were assigned to parcels using MNI coordinates derived from non-linear registration if they fell within 2 mm of a parcel; unassigned samples were discarded, and samples were not mirrored across hemispheres. Expression values were normalized across genes within each tissue sample, and then across tissue samples within each gene, using a scaled robust sigmoid function [102]. Samples assigned to the same parcel were averaged within each donor and then across donors, yielding a matrix ***X****_gene_* of *N* genes (rows) by *P* brain regions (columns). From ***X****_gene_* we retained a curated panel of 2,031 brain-expressed genes previously used to study timescales [14], restricted to 347 of 400 Schaefer parcels that had AHBA coverage.

We decomposed ***X****_gene_* using principal components analysis (PCA). We retained the first three components (PC1–PC3; 18.9%, 11.6%, 9.0% of variance) and report the two that were associated with developmental change (PC1 and PC3) in the main text. To relate each component to development, we correlated its regional score map (Pearson) with the regional rate of change in *|*Δtimescale*|*—the first derivative of the deflection GAM (Fig. 3A)—at each of the 200 evaluated ages. This process gave an age-resolved coupling curve per component (Fig. 5B). Significance was assessed against a spatial null. For each component we generated 10,000 BrainSMASH surrogates of its score map and recomputed the correlations at each age. This process created age-specific empirical null distributions that corresponding age-specific *p*-values were derived from.

To characterize each component, we identified the genes with the strongest positive and negative loadings at each pole, retaining only those that were reproducible under resampling. Specifically, we refit the PCA on 500 bootstrap resamples of parcels. Within each resample, every component of the full-sample (reference) solution was matched to the bootstrap component with which its loading vector shared the highest absolute correlation, and that bootstrap component was sign-flipped to align with the reference. For each reference component, we then counted how often each gene re-entered the top 40 loadings of its pole across resamples. A gene was deemed stable, and thus eligible for retention, if it did so in at least 50% of resamples; the 20 most strongly loading eligible genes were shown at each pole. Under this criterion, PC1 loadings were near-perfectly reproducible (median gene stability 100%), whereas PC2 and PC3 were progressively less so (medians of 84% and 76%). We note that loadings formed a long, shallow tail—by participation ratio, roughly 1,400 of the 2,031 genes contributed substantially—so the displayed genes represent a stable but arbitrary cut, and their rank order within a pole should not be interpreted.

### 4.11 Intracortical myelin

As an *in vivo* proxy for intracortical myelin, we used the ratio of T1-weighted to T2-weighted image intensity (T1w/T2w), which is sensitive to cortical myelin content [103]. For each HCP-D participant, we took the MyelinMap_BC_MSMAll T1w/T2w surface map generated by the HCP minimal preprocessing pipelines [86], and averaged vertex values within each Schaefer-400 parcel. We then averaged these parcellated maps across all participants in our HCP-D sample for whom a T1w/T2w map was available, yielding a single group-level map of intracortical myelin. We related this map to the regional rate of developmental change in timescale deflections using the same age-resolved procedure as for the gene-expression components (Section 4.10), with significance assessed against 10,000 BrainSMASH surrogates of the myelin map.

## Acknowledgements

The authors thank Ahmad Beyh and Emily Heilner for thoughtful discussions. LP was supported by the National Institute Of Mental Health of the National Institutes of Health under Award Number R00MH127296. VJS was supported by the National Institute Of Mental Health of the National Institutes of Health under Award Numbers T32MH018951 and K99MH146391. AW was supported by the National Institute Of Mental Health of the National Institutes of Health under Award Number R00MH125021. BL was supported by the National Institute Of Mental Health of the National Institutes of Health under Award Number R00MH127293.

## Data availability

This study made use of multiple datasets. The Human Connectome Project Young Adult (HCP-YA) sample is available at https://balsa.wustl.edu/. The Welsh Advanced Neuroimaging Database (WAND) is available at https://doi.gin.g-node.org/10.12751/g-node.5mv3bf/. The Human Connectome Project Development (HCP-D) sample is available at https://www.humanconnectome.org/study/hcp-lifespan-development/. The Penn Longitudinal Executive functioning in Adolescent Development (Penn LEAD) dataset is available at https://openneuro.org/datasets/ds007116/. Finally, human gene expression data are available at https://portal.brain-map.org/. Data were provided [in part] by the Human Connectome Project, WU-Minn Consortium (Principal Investigators: David Van Essen and Kamil Ugurbil; 1U54MH091657) funded by the 16 NIH Institutes and Centers that support the NIH Blueprint for Neuroscience Research; and by the McDonnell Center for Systems Neuroscience at Washington University.

## Code availability

Code for this project can be found at https://github.com/LindenParkesLab/intrinsic_neural_timescales under v0.1.0 (preprint release). For the analyses described herein, we used coding assistance from a large language model (Claude Opus v5.5, Anthropic). We manually reviewed and modified all the code used in the analyses and assume full responsibility for the results.

## 5 Supplementary Materials

### Supplementary Figures

**Figure S1.**
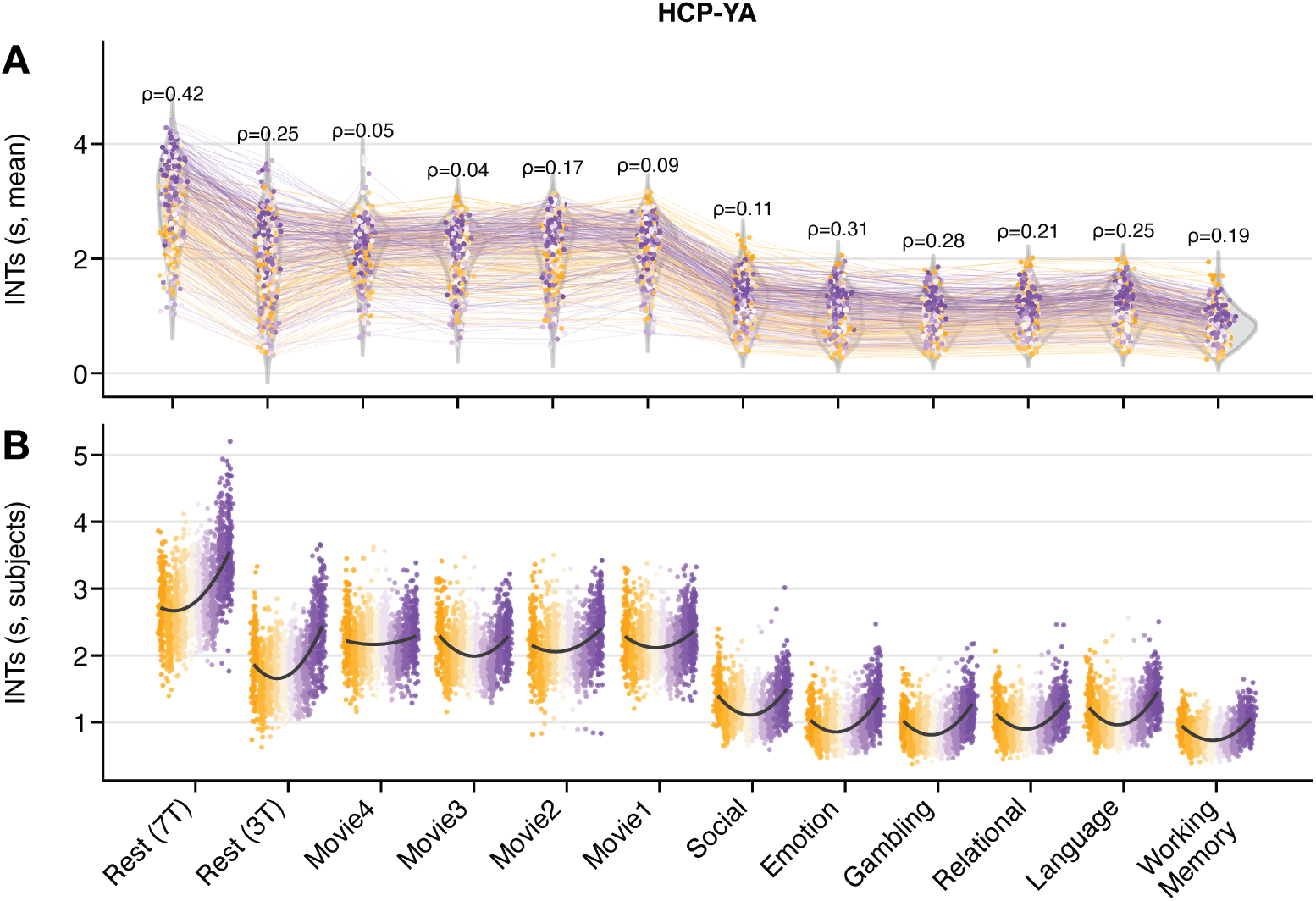
Timescales across individual HCP-YA scans. Here, we show the same HCP-YA data as in Fig. 1, except we display every task and movie separately. (**A**) Distributions of regional timescales. Each point is the timescale for a single brain region averaged over subjects and colored by the S-A axis (orange = sensorimotor pole; purple = association pole). Lines connect the same region across scans, and annotated *ρ* values indicate the Spearman correlation between timescales and the S-A axis. (**B**) Subject-level timescale distributions grouped by S-A axis decile. Here, each point is a timescale from a single subject averaged over regions within a single decile, and the black curve traces the group-mean trend across deciles. These data show that timescales reduced progressively across the movie and task runs relative to rest, with the working memory (WM) task yielding the lowest timescales, and thus the largest deflections from rest.

**Figure S2.**
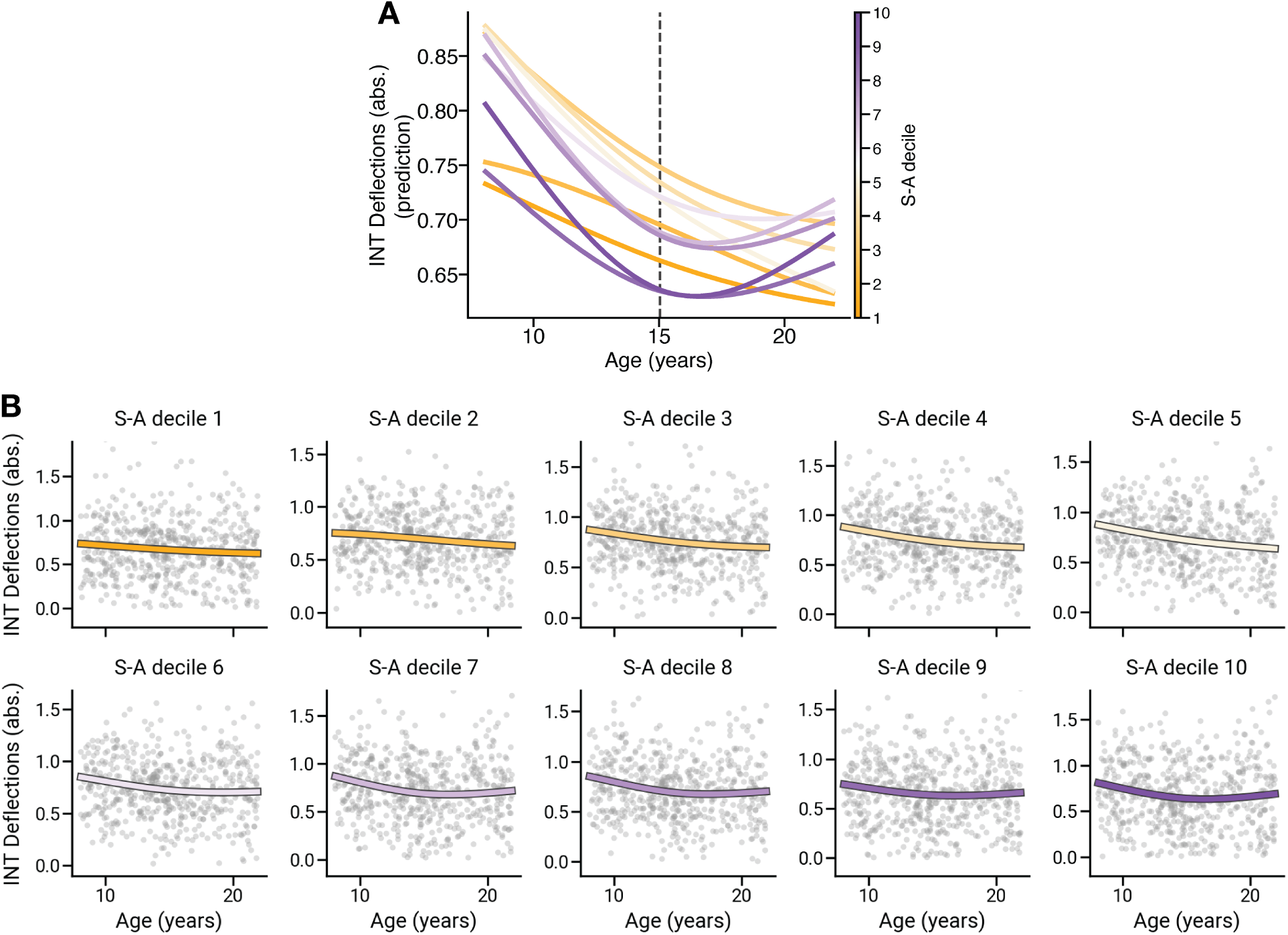
Developmental GAM fits of timescale deflections by S-A decile, without mean-centering. This figure is a companion to Fig. 3B, and shows the same GAM trajectories, for each S-A decile, without mean-centering. (**A**) GAM-predicted absolute timescale deflections for all ten S-A axis deciles (orange = sensorimotor pole; purple = association pole). The dashed vertical line marks the data-driven stage boundary at age 15. (**B**) The same GAM fits shown separately for each S-A decile, each overlaid on subject-level timescale deflection data (grey points; one point per subject).

**Figure S3.**
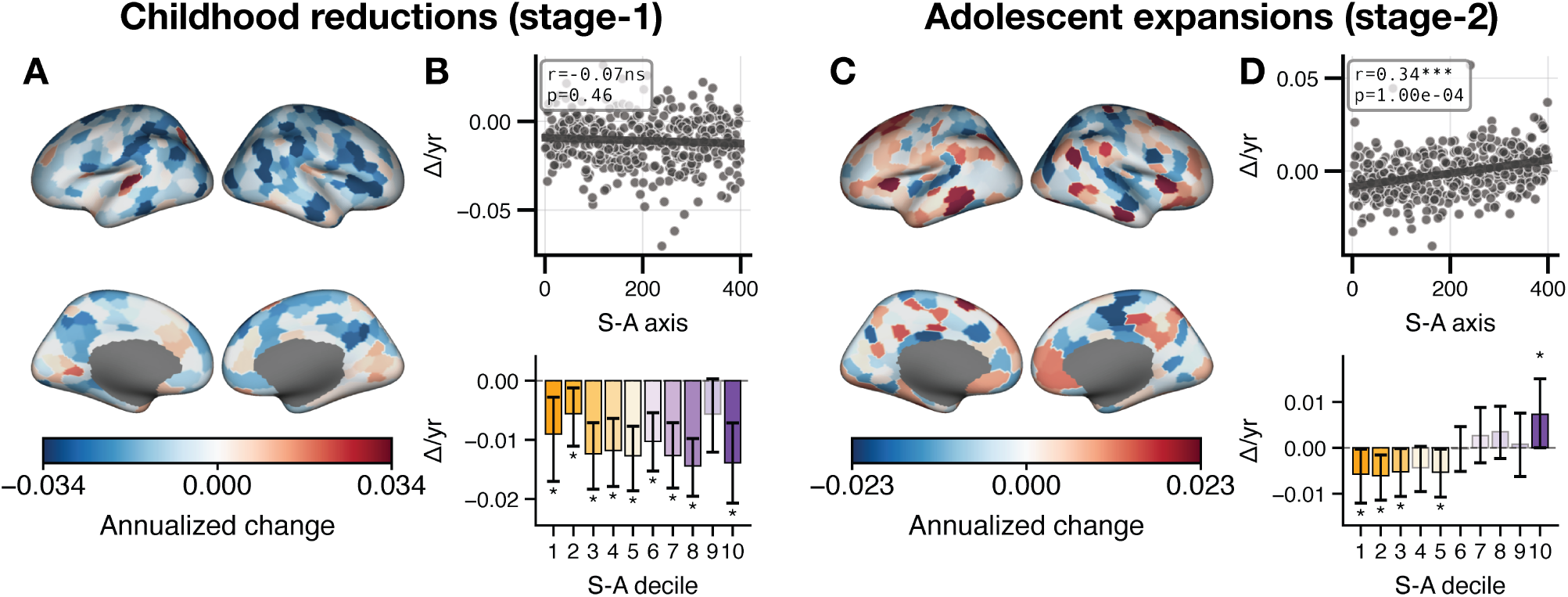
Summary of the two-stage developmental course of timescale deflections. We summarized each developmental stage as a single cortical map by integrating the fitted regional GAM derivatives within the corresponding age window: a stage-1 map spanning childhood (ages 8–15 years) and a stage-2 map spanning adolescence (ages 15–22), each expressed as an annualized rate of change. We then examined whether each stage was organized along the S-A axis. Decile means were tested against zero using a subject bootstrap (5,000 resamples); asterisks mark deciles whose 95% confidence interval excluded zero. (**A**) Stage-1 map of childhood reductions in timescale deflections. (**B**) Stage-1 map as a function of the S-A axis (top) and averaged within S-A deciles (bottom). The stage-1 map was negative across most of the cortex (78.8% of parcels) and reliably negative in nine of the ten S-A deciles, consistent with a generalized reduction in timescale deflections throughout childhood, and showed no relationship to the cortical hierarchy (*r* = *−*0.07, *p*smash = 0.46). Thus, stage 1 captured a global, hierarchy-independent reduction in timescale deflections. (**C**) Stage-2 map of adolescent expansions in timescale deflections. (**D**) Stage-2 map as a function of the S-A axis (top) and averaged within S-A deciles (bottom). The stage-2 map was spatially heterogeneous and correlated positively with the S-A axis (*r* = 0.34, *p*smash *<* 1 *×*10*^−^*^4^). Analysis of the S-A deciles revealed that this positive correlation arose because sensorimotor and mid-hierarchy cortex continued to reduce—deciles 1–3 and 5 reduced significantly—whereas the association pole expanded, with decile 10 showing a reliable net increase. Deciles 4 and 6–9 showed no reliable net change. Thus, stage 2 captured a hierarchically organized pattern of divergence, characterized by a persistence of the childhood reduction at the sensorimotor pole paired with expansion at the association pole.

**Figure S4.**
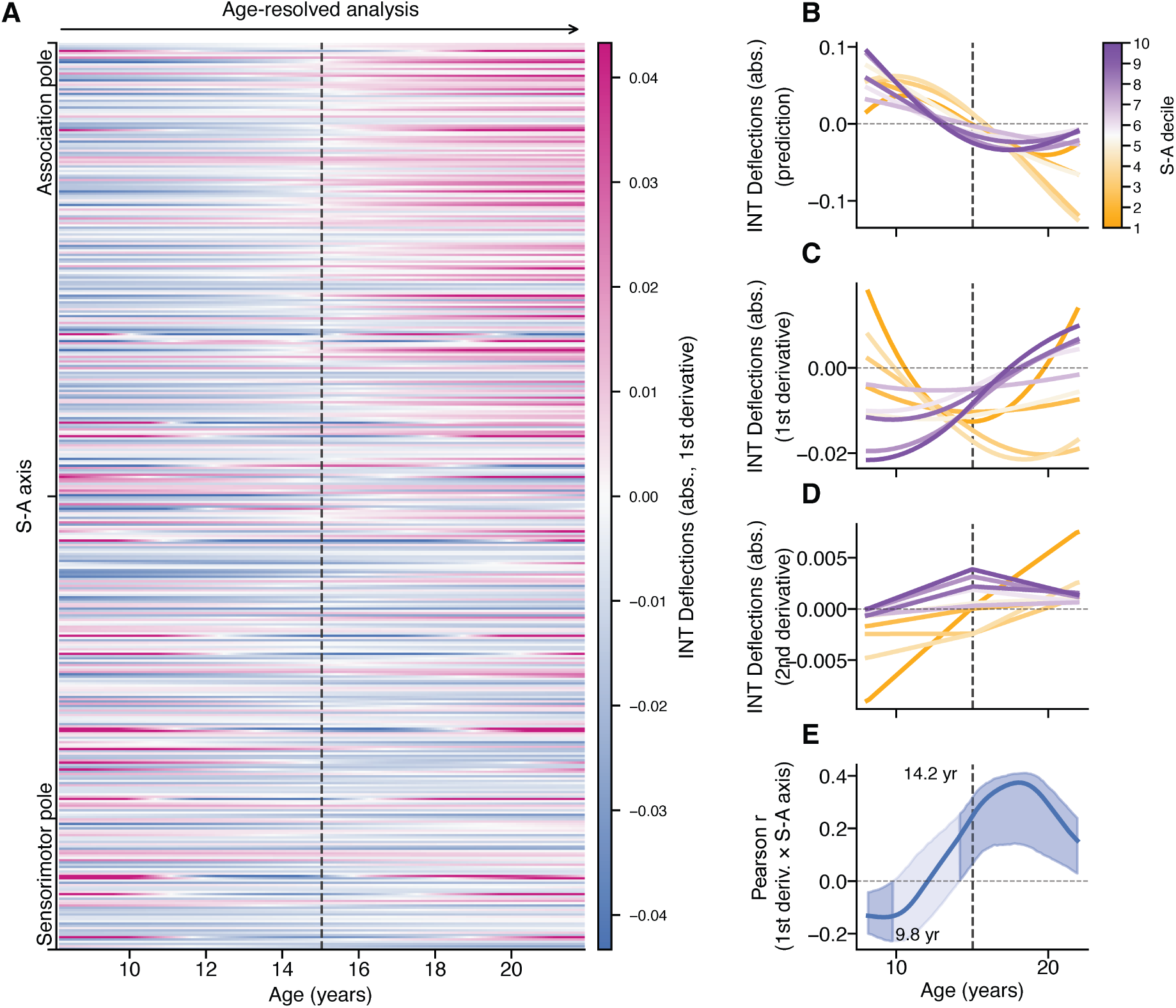
Two-stage developmental course of timescale deflections for the HCP-D Emotion task. This figure is a companion to Fig. 3, but instead shows timescale deflections derived from the Emotion task. (**A**) Heatmap of the annualized rate of change (first derivative of GAM fits) in timescale deflections across development for each cortical region, ordered by the S-A axis (association pole at top, sensorimotor pole at bottom). (**B**) GAM-fitted developmental trajectories for each S-A axis decile (orange = sensorimotor; purple = association), mean-centered. (**C**) First temporal derivatives of the trajectories in **B**. (**D**) Second derivatives, which provide acceleration per decile. (**E**) Pearson correlation between regional rates of change in timescale deflections and the S-A axis, computed at monthly intervals, with the subject-level bootstrap 95% CI (shaded band). This reproduces the two-stage course reported in the main text: a childhood reduction giving way to hierarchically organized expansion in association cortex.

**Figure S5.**
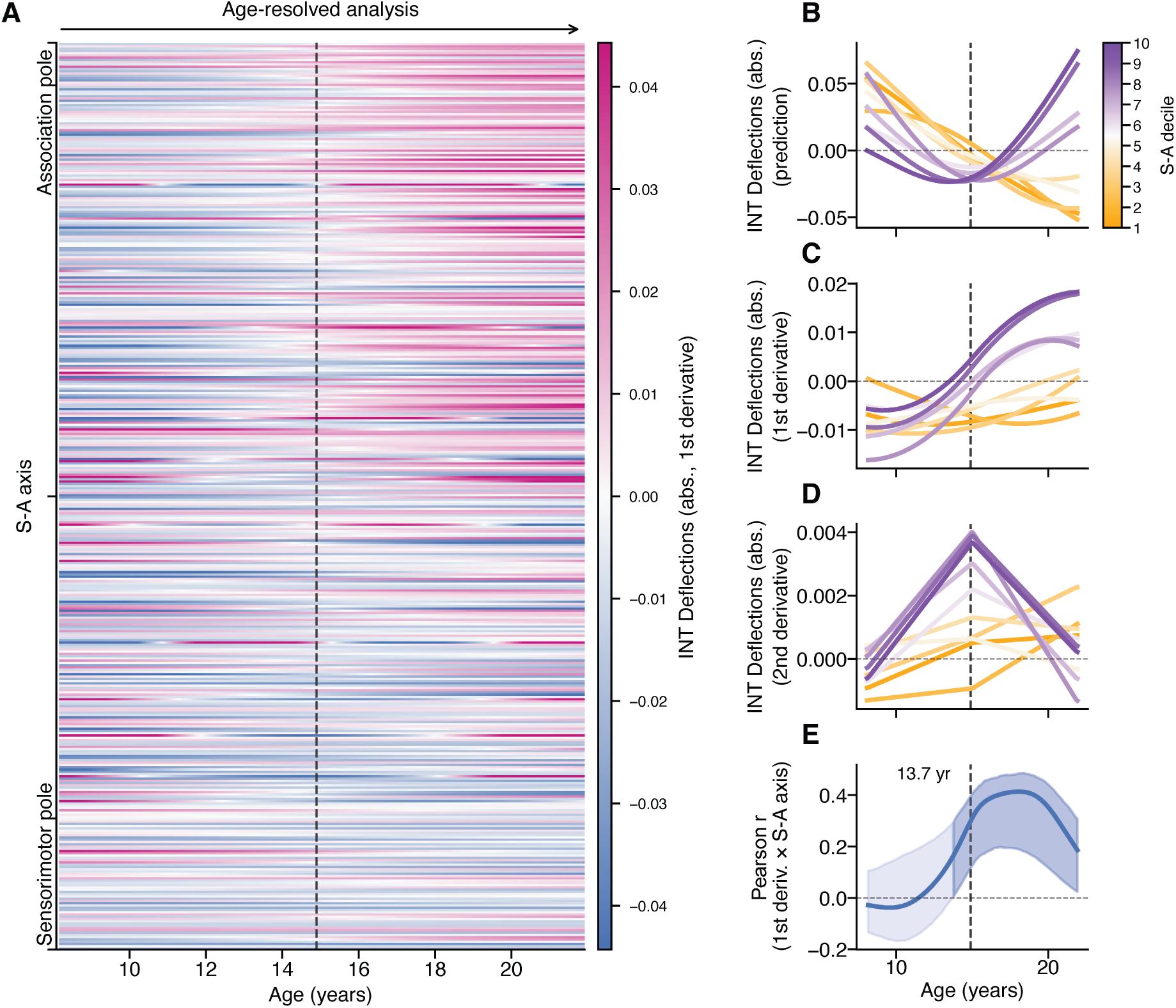
Two-stage developmental course of timescale deflections for the HCP-D Guessing task. This figure is a companion to Fig. 3, but instead shows timescale deflections derived from the Guessing task. (**A**) Heatmap of the annualized rate of change (first derivative of GAM fits) in timescale deflections across development for each cortical region, ordered by the S-A axis (association pole at top, sensorimotor pole at bottom). (**B**) GAM-fitted developmental trajectories for each S-A axis decile (orange = sensorimotor; purple = association), mean-centered. (**C**) First temporal derivatives of the trajectories in **B**. (**D**) Second derivatives, which provide acceleration per decile. (**E**) Pearson correlation between regional rates of change in timescale deflections and the S-A axis, computed at monthly intervals, with the subject-level bootstrap 95% CI (shaded band). This reproduces the two-stage course reported in the main text: a childhood reduction giving way to hierarchically organized expansion in association cortex.

**Figure S6.**
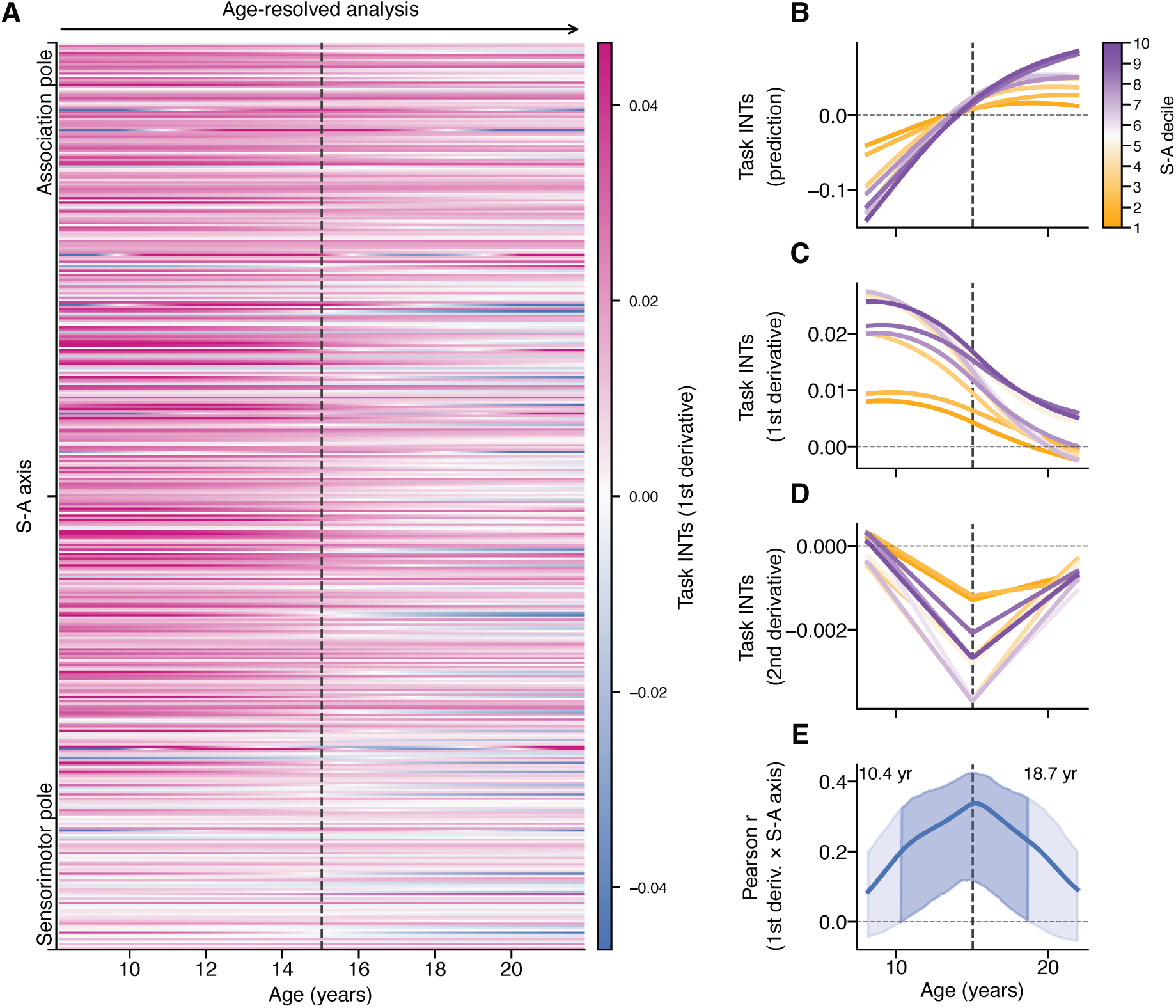
Development of Carit task-based timescales along the cortical hierarchy. This figure is a companion to Fig. 3, but with the dependent variable in all GAMs being task-based timescales rather than absolute timescale deflections. (**A**) Heatmap of the annualized rate of change (first derivative of GAM fits). (**B**) GAM-fitted developmental trajectories for each S-A axis decile (orange = sensorimotor; purple = association), mean-centered. (**C**) First temporal derivatives of the trajectories in **B**. (**D**) Second derivatives, which provide acceleration per decile. (**E**) Pearson correlation between regional rates of change in task-based timescales and the S-A axis, computed at monthly intervals, with the subject-level bootstrap 95% CI (shaded band).

### 5.1 Regression of scan structure

As discussed in the main text, a key methodological decision in our analyses was to regress out the influence of scan structure from the fMRI time series prior to estimating timescales from the autocorrelation function (ACF). For task-based data, this involved removing the influence of the block or event-related structure of the task design. For movie-watching data, this involved removing the movie-clip structure, which resembled a block design. Here, we document the rationale for this decision, as well as its impact.

#### 5.1.1 Rationale

Given the temporally slow nature of fMRI BOLD data, the presence of a block-design or a movie-clip structure necessarily increases the ACF of the underlying time series data, which in turn increases the timescale. This is because the temporal structure of the task is itself autocorrelated. For example, block designs alternate between task conditions that can span tens of seconds, with interleaved periods where no task is presented that may also last several seconds. As such, the corresponding regressors carry most of their power at low frequencies. Additionally, the convolution with the hemodynamic response function extends that dependence further in time. The consequence is that any brain region whose activity tracks this design inherits this dependence. Critically, because we estimate timescales from the decay of the ACF, this inherited structure affects the timescale estimate directly because the ACF is driven by both an intrinsic component and a task-evoked component, and the task-evoked component decays on the timescale of the design rather than that of the region. Fig. S7 shows this effect in a single subject from the HCP-YA.

**Figure S7.**
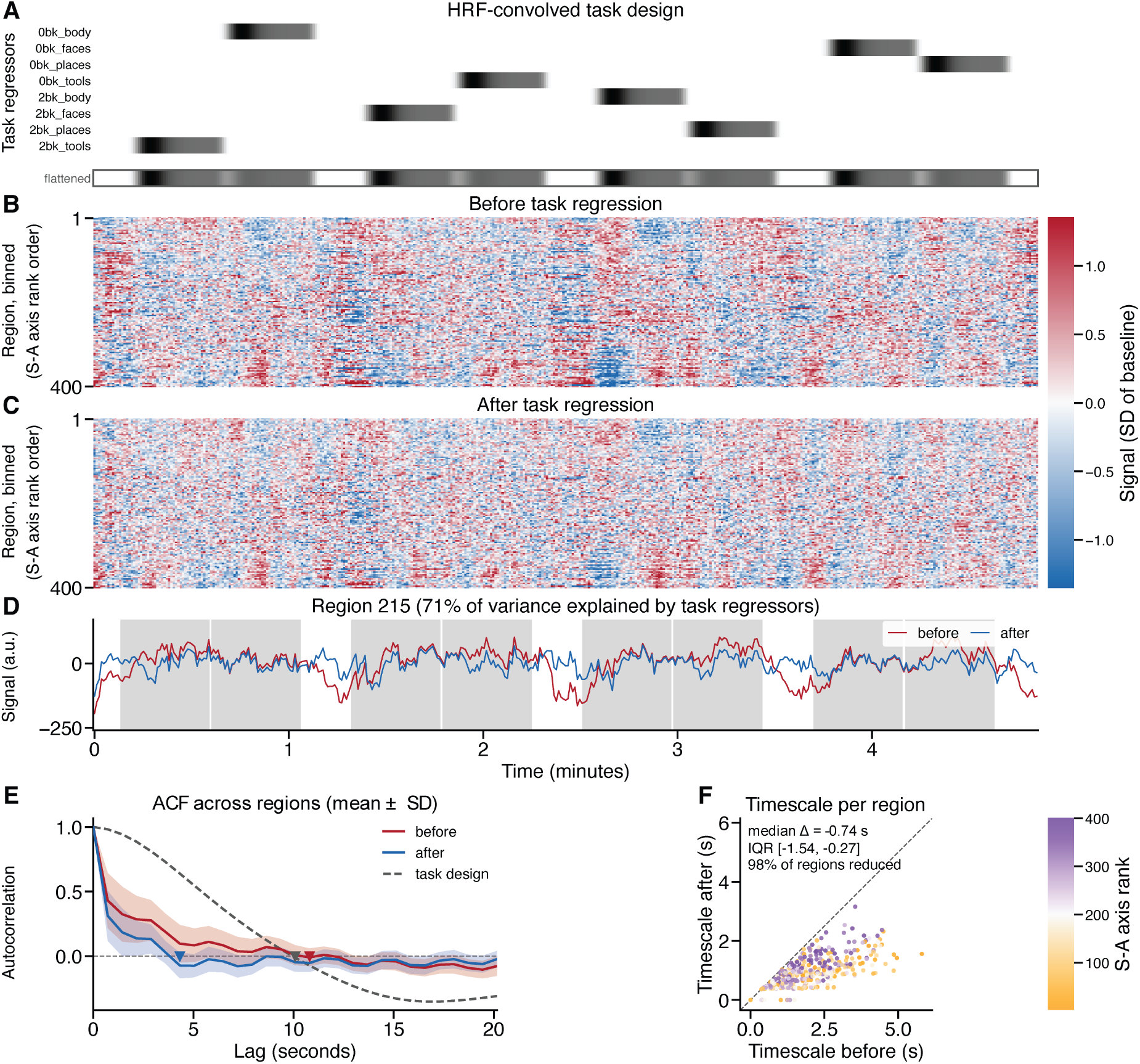
Example time series data for the working memory task (HCP-YA subject 214423) with and without regression of the task design. Here, we show data for a single working memory run (tfMRIWMLR; 405 frames). Throughout this figure, “before” denotes the residual of a general linear model containing only a cosine drift basis and a constant, and “after” denotes the residual of the full model that also contained the task regressors. (**A**) The HRF-convolved task design, with one regressor per condition, and beneath it the flattened regressor obtained by summing across conditions (outlined). The design comprises eight 27.5 second blocks arranged as back-to-back pairs separated by fixation, giving a repeating cycle of 71.3 seconds. (**B, C**) Parcellated time series before and after regression of the design, for all 400 parcels ordered along the S-A axis. For display only in this figure, we scaled each parcel by the standard deviation of its own time series before task regression, and applied the identical scaling in both panels so that the variance removed by regression remained visible. (**D**) An example single parcel wherein variance was most reduced by regression of the task design (region 215; 71% explained). Task blocks are shaded. The time series before task regression (red) visibly tracks the block structure, whereas the residual after task regression (blue) does not. (**E**) ACF averaged across parcels (mean *±* SD) before (red) and after (blue) regression, shown against the ACF of the task design itself (dashed line). The task design ACF was estimated from the flattened version shown in panel A. The ACF for the task design remained positive out to 10 seconds, reached a minimum at 54.0 seconds (not shown), and returned to a local maximum at 71.3 seconds (not shown), which was the period of the block structure. That is, the design itself was strongly autocorrelated on the timescale of the scan. Accordingly, the parcel-averaged ACF crosses zero at 10.8 seconds before regression (red arrowhead) but at 4.3 seconds after (blue arrowhead). (**F**) Timescales for each parcel before versus after task regression. Color denotes S-A axis rank and the dashed line is the identity. Timescales were reduced in 98% of parcels (mean 1.90 seconds to 0.93 seconds; median change *−*0.74 seconds).

Since our goal was to examine how cognitive tasks (and movies) impacted the intrinsic component and not the task-evoked component, we chose to regress out scan structure from all task and movie watching scans. Nevertheless, below we present a series of supplementary figures that illustrate how this choice impacted our results. See Section 5.1.2 below.

#### 5.1.2 Impact

Below, we show a set of plots comprising several of our key analyses without regression of scan structure. Fig. S8 illustrates timescales from all scans for the HCP-YA without scan regression. Fig. S9 and Fig. S10 show behavioral correlations in HCP-YA (cf. Fig. 2) without scan regression. The magnitudes of the brain–behavior correlations were unchanged by regression, but their organization along the S-A axis was not preserved.

**Figure S8.**
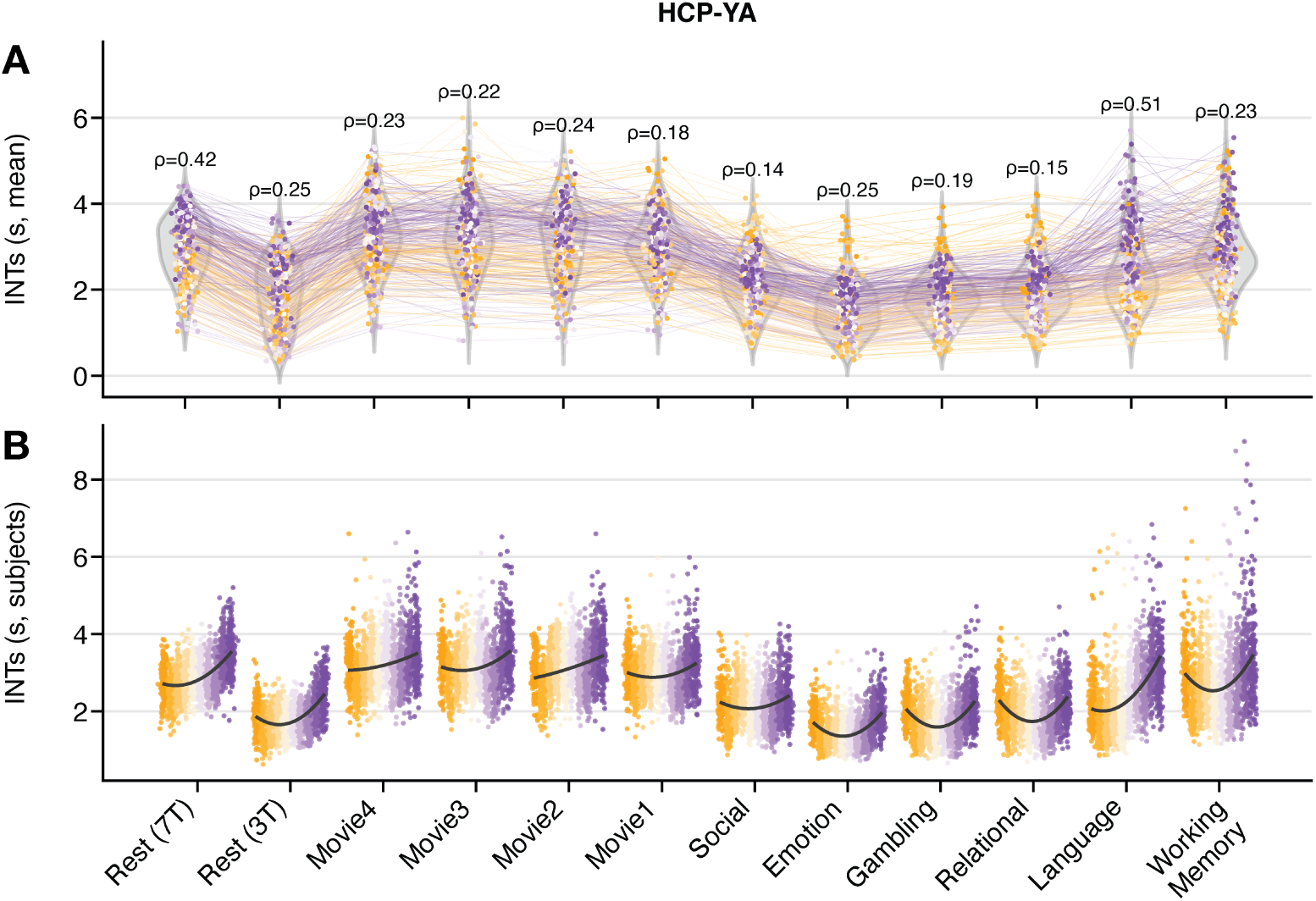
HCP-YA timescales across all scans without regression of scan design. Here, we show the same HCP-YA data as in Fig. S1, but with timescales computed from the raw regional time series without first regressing out the block-design, event-related, or movie-clip scan structure. (**A**) Distributions of regional timescales. (**B**) Subject-level timescale distributions grouped by S-A axis decile.

**Figure S9.**
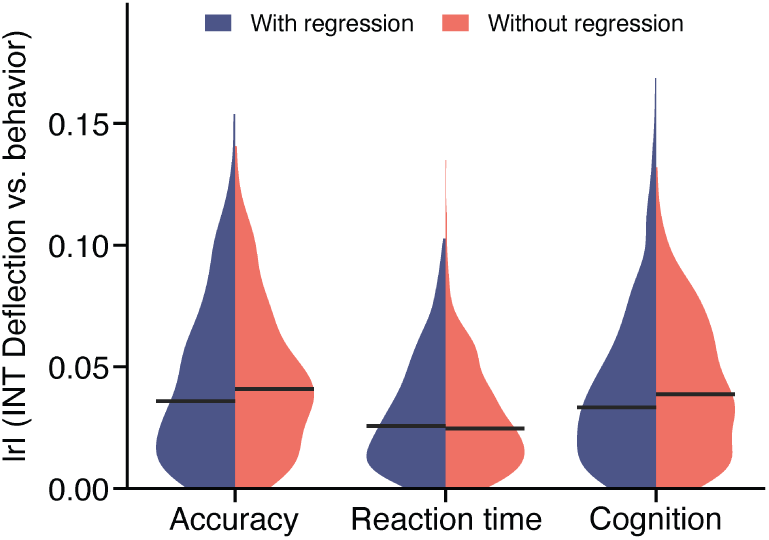
Correlations between timescale deflections and behavior with and without regression of scan design. Here, we show the absolute correlations from Fig. 2C against the same effects estimated without regressing the task design from the working memory (WM) time series. No differences in effect size magnitudes were observed. Thus, regression of scan design had no impact on the size of brain-behavior correlations in the HCP-YA WM task.

**Figure S10.**
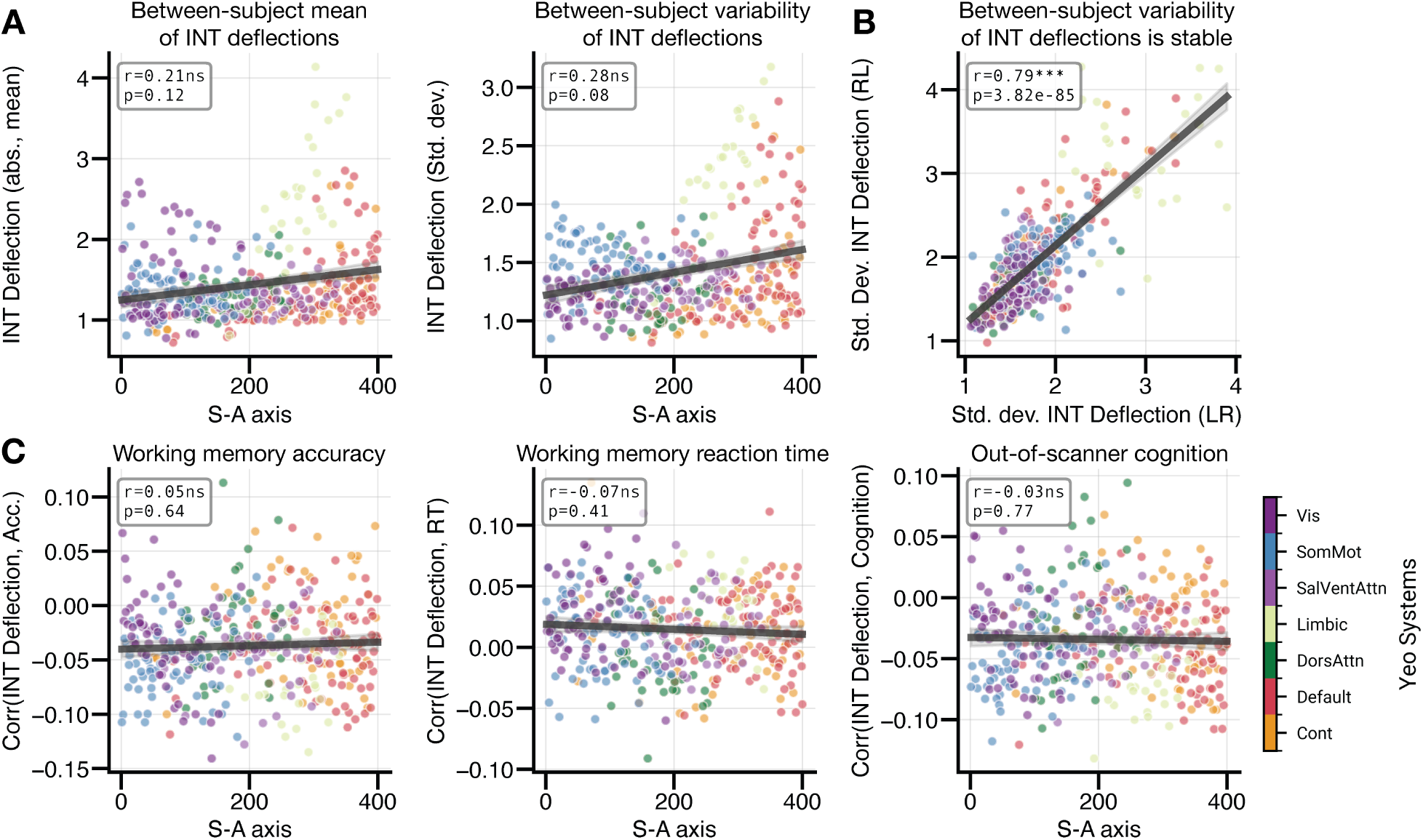
Spatial organization and behavioral relevance of timescale deflections without regression of scan design. Here, we show the same HCP-YA data as in Fig. 2, but computed from timescale deflections derived without regressing the task design from the WM time series (cf. Fig. S8). (**A**) Between-subject mean (left) and standard deviation (right) of absolute timescale deflections against S-A axis rank; each dot is one region colored by Yeo network. Neither the mean nor the variability was correlated significantly with the S-A axis (*r* = 0.21, *p*smash = 0.12; *r* = 0.28, *p*smash = 0.08). (**B**) Between-subject variability estimated from the LR versus RL scan acquisitions. (**C**) Region-wise correlations between absolute timescale deflections and working memory accuracy (left), reaction time (middle), and out-of-scanner cognition (right), plotted against the S-A axis. None were organized along the cortical hierarchy (*r* = 0.05, *p*smash = 0.64; *r* = *−*0.07, *p*smash = 0.41; *r* = *−*0.03, *p*smash = 0.77). Thus, without regression of scan design, the correlations between timescale deflections and behavior were no longer hierarchically organized, indicating that these effects depend on isolating intrinsic dynamics from stimulus-locked variance.

#### 5.1.3 Regression of scan structure applied to resting-state data

As noted above and in the main text, timescales for task scans were estimated from the residuals of a GLM designed to remove scan structure, while timescales for rest were estimated without that GLM. Importantly, projecting a task design out of our fMRI timeseries removes low-frequency variance, and because the timescale is a product of the decay of the ACF, that projection shortens the estimate (as we demonstrate above). As such, part of the task–rest difference that we call *timescale deflections* (Figs. 1–5) may be a property of the processing rather than of the various task conditions, and our inferences rely on assuming an effect of those conditions.

Here, we conduct a sensitivity analysis designed to address this issue by regressing scan structure out of the resting-state data in the HCP-YA and HCP-D datasets. Specifically, for each task run we constructed a matched control GLM by fitting that task run’s design matrix to a resting-state run from the same participant, matched on acquisition, phase-encoding direction and scan length. Scan length was matched by truncating resting-state runs to the length of the task run, as in the main analyses of Figs. 2–5 (see Section 4.4), so this control differs from those analyses only in that the task design was additionally projected out of the resting-state data. Critically, this control contains no task-related variance in the underlying data, so any reduction in resting-state timescales following regression is solely a product of processing, and any difference in timescales between a task residual and its own resting-state control is what survives this processing. Applying this control, we found that timescale deflections remained negative in every condition across both datasets (Fig. S11), though their magnitude did diminish. These results demonstrate that the reduction in timescales during task compared to rest is not an artifact of the regression of scan structure.

**Figure S11.**
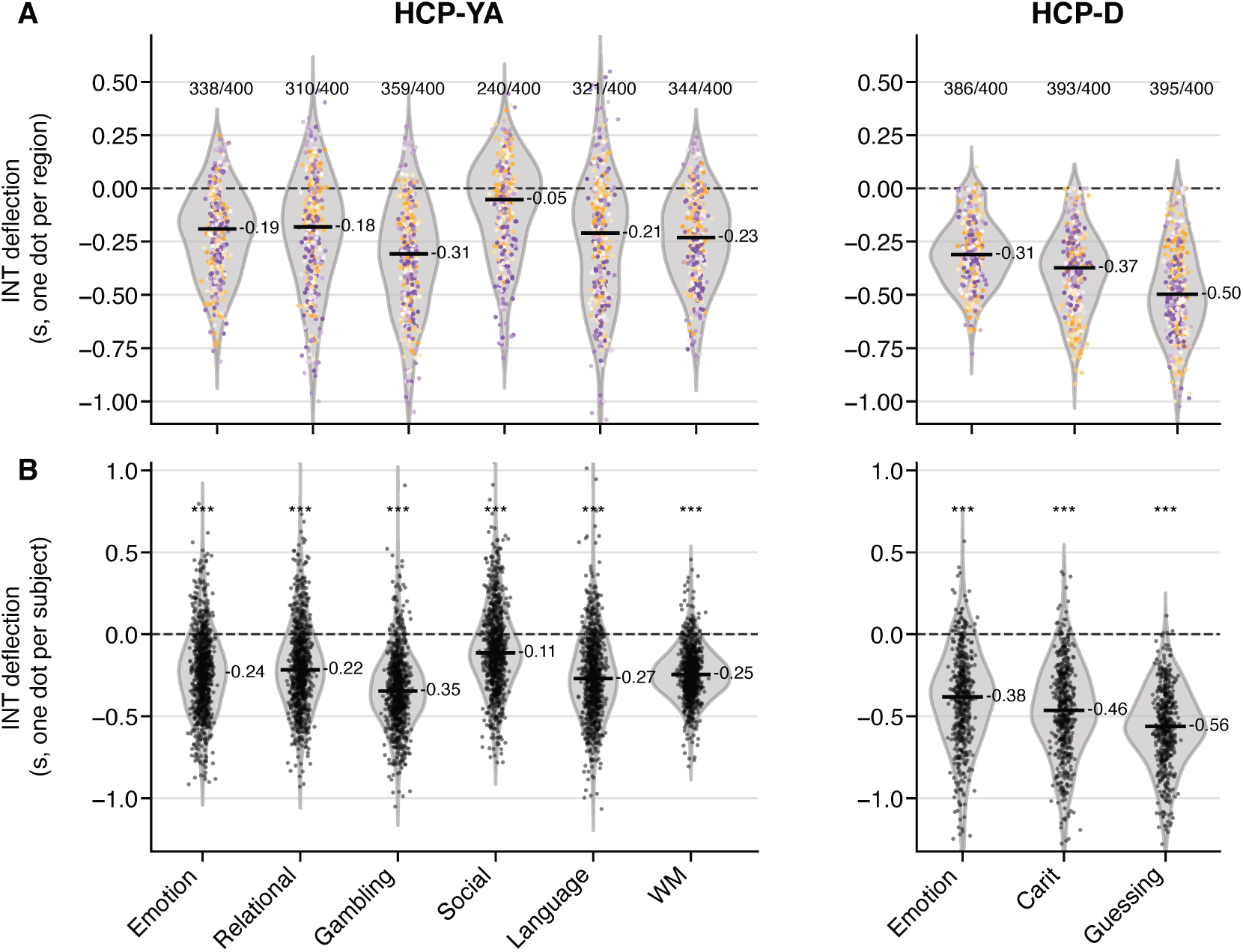
Timescale deflections survive a control regression wherein regression of scan structure was applied to resting-state data. For every task run, we built a matched control GLM by fitting that run’s design matrix to a resting-state run from the same participant, matched on acquisition, phase-encoding direction and scan length. Here, we show timescale deflections computed using the task data used in the main text minus its resting-state data that underwent our matched control. (**A**) Timescale deflections for each brain region, averaged over subjects and colored by S-A axis rank. The annotations show the number of parcels with a negative deflection. (**B**) Timescale deflections for each subject, averaged over regions. Asterisks denote a two-sided one-sample Wilcoxon test against zero, FDR-corrected across conditions (*pFDR <* 0.001). In both plots, horizontal black bars mark the median deflections. Deflections are negative in every condition: HCP-YA *−*0.11 to *−*0.35 s, HCP-D *−*0.38 to *−*0.56 s across participants. The Social task from HCP-YA was the weakest case, reliable across participants (*−*0.11 s) but negative in only 240 of 400 parcels.

### Boilerplate material

#### 5.1.4 fMRIprep

Results included in this manuscript come from preprocessing performed using *fMRIPrep* 24.1.1 (Esteban et al. [104]; Esteban et al. [105]; RRID:SCR_016216), which is based on *Nipype* 1.8.6 (Gorgolewski et al. [106]; Gorgolewski et al. [107]; RRID:SCR_002502).

##### Anatomical data preprocessin

A total of 1 T1-weighted (T1w) images were found within the input BIDS dataset. The T1w image was corrected for intensity non-uniformity (INU) with N4BiasFieldCorrection [108], distributed with ANTs 2.5.3 [109, RRID:SCR_004757], and used as T1w-reference throughout the workflow. The T1w-reference was then skull-stripped with a *Nipype* implementation of the antsBrainExtraction.sh workflow (from ANTs), using OASIS30ANTs as target template. Brain tissue segmentation of cerebrospinal fluid (CSF), white-matter (WM) and gray-matter (GM) was performed on the brain-extracted T1w using fast [FSL (version unknown), RRID:SCR_002823, 110]. Brain surfaces were reconstructed using recon-all [FreeSurfer 7.3.2, RRID:SCR_001847, 111], and the brain mask estimated previously was refined with a custom variation of the method to reconcile ANTs-derived and FreeSurfer-derived segmentations of the cortical gray-matter of Mindboggle [RRID:SCR_002438, 112]. A T2-weighted image was used to improve pial surface refinement. Brain surfaces were reconstructed using recon-all [FreeSurfer 7.3.2, RRID:SCR_001847, 111], and the brain mask estimated previously was refined with a custom variation of the method to reconcile ANTs-derived and FreeSurfer-derived segmentations of the cortical gray-matter of Mindboggle [RRID:SCR_002438, 112]. Volume-based spatial normalization to one standard space (MNI152NLin2009cAsym) was performed through nonlinear registration with antsRegistration (ANTs 2.5.3), using brain-extracted versions of both T1w reference and the T1w template. The following template was were selected for spatial normalization and accessed with *TemplateFlow* [24.2.0, 113]: *ICBM 152 Nonlinear Asymmetrical template version 2009c* [Fonov et al. [114], RRID:SCR_008796; TemplateFlow ID: MNI152NLin2009cAsym].

##### Functional data preprocessing

For each of the 2 BOLD runs found per subject (across all tasks and sessions), the following preprocessing was performed. First, a reference volume was generated, using a custom methodology of *fMRIPrep*, for use in head motion correction. Head-motion parameters with respect to the BOLD reference (transformation matrices, and six corresponding rotation and translation parameters) are estimated before any spatiotemporal filtering using mcflirt [FSL, 115]. The BOLD reference was then co-registered to the T1w reference using bbregister (FreeSurfer) which implements boundary-based registration [116]. Co-registration was configured with six degrees of freedom. The aligned T2w image was used for initial co-registration.Several confounding time-series were calculated based on the *preprocessed BOLD*: framewise displacement (FD), DVARS and three region-wise global signals. FD was computed using two formulations following Power (absolute sum of relative motions, Power et al. [117]) and Jenkinson (relative root mean square displacement between affines, Jenkinson et al. [115]). FD and DVARS are calculated for each functional run, both using their implementations in *Nipype* [following the definitions by 117]. The three global signals are extracted within the CSF, the WM, and the whole-brain masks. Additionally, a set of physiological regressors were extracted to allow for component-based noise correction [*CompCor*, 118]. Principal components are estimated after high-pass filtering the *preprocessed BOLD* time-series (using a discrete cosine filter with 128s cut-off) for the two *CompCor* variants: temporal (tCompCor) and anatomical (aCompCor). tCompCor components are then calculated from the top 2% variable voxels within the brain mask. For aCompCor, three probabilistic masks (CSF, WM and combined CSF+WM) are generated in anatomical space. The implementation differs from that of Behzadi et al. in that instead of eroding the masks by 2 pixels on BOLD space, a mask of pixels that likely contain a volume fraction of GM is subtracted from the aCompCor masks. This mask is obtained by dilating a GM mask extracted from the FreeSurfer’s *aseg* segmentation, and it ensures components are not extracted from voxels containing a minimal fraction of GM. Finally, these masks are resampled into BOLD space and binarized by thresholding at 0.99 (as in the original implementation). Components are also calculated separately within the WM and CSF masks. For each CompCor decomposition, the *k* components with the largest singular values are retained, such that the retained components’ time series are sufficient to explain 50 percent of variance across the nuisance mask (CSF, WM, combined, or temporal). The remaining components are dropped from consideration. The head-motion estimates calculated in the correction step were also placed within the corresponding confounds file. The confound time series derived from head motion estimates and global signals were expanded with the inclusion of temporal derivatives and quadratic terms for each [119]. Frames that exceeded a threshold of 0.5 mm FD or 1.5 standardized DVARS were annotated as motion outliers. Additional nuisance timeseries are calculated by means of principal components analysis of the signal found within a thin band (*crown*) of voxels around the edge of the brain, as proposed by [120]. All resamplings can be performed with *a single interpolation step* by composing all the pertinent transformations (i.e. head-motion transform matrices, susceptibility distortion correction when available, and co-registrations to anatomical and output spaces). Gridded (volumetric) resamplings were performed using nitransforms, configured with cubic B-spline interpolation.

Many internal operations of *fMRIPrep* use *Nilearn* 0.10.4 [121, RRID:SCR_001362], mostly within the functional processing workflow. For more details of the pipeline, see the section corresponding to workflows in *fMRIPrep*’s documentation.

#### 5.1.5 Post-processing of fmriprep outputs

The eXtensible Connectivity Pipeline- DCAN (XCP-D) [122–124] was used to post-process the outputs of *fMRIPrep* version 24.1.1 [125, 126, RRID:SCR_016216]. XCP-D was built with *Nipype* version 1.8.6 [106, RRID:SCR_002502].

##### Segmentation

The following atlases were used in the workflow: the Schaefer Supplemented with Subcortical Structures (4S) atlas [127–131] at 3 different resolutions (156, 256, and 456 parcels), the Schaefer1007 atlas, the Schaefer2007 atlas, and the Schaefer4007 atlas. Each atlas was warped to the same space and resolution as the BOLD file.

##### Anatomical dat

Native-space T1w images were transformed to MNI152NLin2009cAsym space at 1 mm3 resolution.

##### Functional dat

For each of the one BOLD runs found per subject (across all tasks and sessions), the following post-processing was performed.

In total, 36 nuisance regressors were selected from the preprocessing confounds, according to the ‘36P’ strategy. These nuisance regressors included six motion parameters, mean global signal, mean white matter signal, mean cerebrospinal fluid signal with their temporal derivatives, and quadratic expansion of six motion parameters, tissue signals and their temporal derivatives [124, 132].

The BOLD data were despiked with *AFNI*’s *3dDespike*.

Nuisance regressors were regressed from the BOLD data using a denoising method based on *Nilearn*’s approach. The timeseries were band-pass filtered using a(n) second-order Butterworth filter, in order to retain signals between 0.01-0.08 Hz. The same filter was applied to the confounds. The resulting time series were then denoised using linear regression. The denoised BOLD was smoothed using *Nilearn* with a Gaussian kernel (FWHM=6 mm).

The amplitude of low-frequency fluctuation (ALFF) [133] was computed by transforming the mean-centered, standard deviation-normalized, denoised BOLD time series to the frequency domain. The power spectrum was computed within the 0.01-0.08 Hz frequency band and the mean square root of the power spectrum was calculated at each voxel to yield voxel-wise ALFF measures. The resulting ALFF values were then multiplied by the standard deviation of the denoised BOLD time series to retain the original scaling. The ALFF maps were smoothed with Nilearn using a Gaussian kernel (FWHM=6 mm). Regional homogeneity (ReHo) [134] was computed with neighborhood voxels using *AFNI*’s *3dReHo* [135].

Processed functional timeseries were extracted from the residual BOLD signal with *Nilearn’s NiftiLabelsMasker* for the atlases. Corresponding pair-wise functional connectivity between all regions was computed for each atlas, which was operationalized as the Pearson’s correlation of each parcel’s unsmoothed timeseries. In cases of partial coverage, uncovered voxels (values of all zeros or NaNs) were either ignored (when the parcel had >50.0% coverage) or were set to zero (when the parcel had <50.0% coverage).

Many internal operations of *XCP-D* use *AFNI* [136, 137], *ANTS* [138], *TemplateFlow* version 24.2.2 [139], *matplotlib* version 3.9.2 [140], *Nibabel* version 5.3.0 [141], *Nilearn* version 0.10.4 [142], *numpy* version 2.1.2 [143], *pybids* version 0.17.2 [144], and *scipy* version 1.14.1 [145]. For more details, see the *XCP-D* website (https://xcp-d.readthedocs.io).

